# The role of plant polyploidy in the structure of plant-pollinator communities

**DOI:** 10.1101/2025.08.04.668489

**Authors:** Keren Halabi, Noa Ecker, Nathália Susin Streher, Tal Pupko, Tia-Lynn Ashman, Itay Mayrose

**Affiliations:** School of Plant Sciences and Food Security, George S. Wise Faculty of Life Sciences, Tel Aviv University, Tel Aviv 69978, Israel; The Shmunis School of Biomedicine and Cancer Research, George S. Wise Faculty of Life Sciences, Tel Aviv University, Tel Aviv 69978, Israel; Department of Biological Sciences, University of Pittsburgh, Pittsburgh, PA 15260, USA

**Keywords:** Evolution, Polyploidy, Plant-pollinator interactions, Networks, Extinction

## Abstract

Polyploidization is a major macromutation, bearing notable genomic and ecological consequences. While the impact of polyploidy on plant abiotic niches is well studied, our understanding of its consequences on biotic interactions, and particularly pollination, is lacking and hardly considers its role in shaping community structure. Here, we integrate hundreds of plant-pollinator networks, ploidy inferences, reproductive traits, and climatic attributes to ascertain whether a general pattern characterizes the link between polyploid frequency and community structure. We further examine whether environmental factors and plant traits known to be associated with polyploidy mediate this relationship. Our analysis reveals that an increased frequency of polyploid species is positively associated with network nestedness while being negatively associated modularity. Structural equation modeling reveals that these associations are partially mediated via the frequency of self-compatible plants and to a lesser extent by differences in flower shape. Despite these alterations in community structure, the heightened abundance of polyploids appears to have minimal impact on network connectance and resilience to extinction. Our findings imply that unlike abiotic interactions, the impact of polyploidy on biotic interactions is less predictable, and is affected synergistically by both phenotypic and environmental factors. Collectively, however, polyploidy still exerts an influence on community structure.

## Introduction

To date, the role of evolution in shaping ecological communities remains largely unexplored (Weber *et al*., 2017; Segar *et al*., 2020). Polyploidy, the possession of more than two set of chromosomes resulting from whole genome duplication (WGD) (Soltis *et al*., 2009; Van De Peer *et al*., 2017; Baduel *et al*., 2018), is a prominent evolutionary process in plants. Most plant species have experienced WGD at some point in their history (Cui *et al*., 2006; Jiao *et al*., 2011), and recent polyploidization events are estimated to have occurred in approximately 35% of extant flowering plant species (Wood *et al*., 2009; Halabi *et al*., 2023). Polyploidy has far reaching genomic consequences (Soltis & Soltis, 2000; Otto, 2007), contributing to the emergence of novel genetic variation that facilitates ecological diversification and rapid niche differentiation (Levin, 1983; Ramsey & Schemske, 2003; Martin & Husband, 2009; Van De Peer *et al*., 2017; Baniaga *et al*., 2020), and potentially affects community assembly and stability (Bergamo *et al*., 2020). Nevertheless, our understanding of the ecological consequences of polyploidy remains limited (Ramsey & Ramsey, 2014; Segraves, 2017).

To date, research on the ecological effects of polyploidy has mostly centered on the abiotic niche (Brittingham *et al*., 2018; Baduel *et al*., 2018; Rice *et al*., 2019; Wei *et al*., 2019; Baniaga *et al*., 2020). Broad-scale comparative analyses have revealed, for example, that polyploids tend to exhibit elevated rates of climatic niche differentiation from their parental species compared to diploids (Baniaga *et al*., 2020), and that polyploids are more common in regions characterized by cold climates and lower species richness (Rice *et al*., 2019). Yet, we lack a similar panoramic view of the effect of polyploidy on biotic interactions that are crucial for community dynamics (Segraves & Anneberg, 2016; Segraves, 2017; Gaynor et al., 2018; Streher et al., 2023; Anneberg et al., 2023) and consequently, we have not assessed whether polyploids contribute uniquely to ecological communities.

Animal-mediated pollination is a biotic interaction essential for the successful reproduction of the majority of flowering plant species (Ollerton et al., 2011; Rodger et al., 2021), and has a significant impact on plant population dynamics (Ashman *et al*., 2004) and community structure (Sargent & Ackerly, 2008; Wei *et al*., 2021; Bascompte & Scheffer, 2022). Changes in climate have been suggested to reduce the availability of pollinators within communities (Doré *et al*., 2021) and could consequently increase competition for pollinators (Mustajärvi *et al*., 2001; Lundgren *et al*., 2016) and jeopardize the resilience of plant-pollinator communities (Johnson *et al*., 2022). However, increased self-fertilization, that has been associated with several plant traits (Rodger et al., 2021), could lessen these consequences. Polyploidy has also been linked with multiple plant traits, including flower size, floral morphology, and self-compatibility (Ramsey & Schemske, 2003), all of which are known to influence pollinator interactions. However, its association with self-fertilization capabilities has been debated (Mable, 2004). Given the pivotal role of polyploidy in plant evolution and its association with diverse plant phenotypes, it is expected to influence pollination niche breadth (i.e., the range of pollinator interactions), akin to its effects on climatic niche differentiation (Padilla-García *et al*., 2023). Therefore, understanding how polyploidy reshape network structure may reveal how evolutionary history translates to ecological assembly, resilience, and function.

Emerging polyploid lineages face an initial frequency disadvantage, primarily because a significant portion of their offspring, resulting from mating with diploids, suffer from genomic incompatibilities and are sterile (Ranney, 2006; Köhler *et al*., 2010). The success of a new polyploid individual in accessing mates of its own cytotype and overcoming this frequency disadvantage is heavily influenced by pollination with two possible trajectories. First, changes in morphology or phenology, e.g., due to the *gigas* effects of neopolyploids (Ramsey & Schemske, 2003), may increase the probability of a new polyploid successfully mating via recruitment of new pollinators, thus shifting or broadening its pollination niche (Segraves & Anneberg, 2016; Casazza *et al*., 2017). On the other hand, narrowing the pollination niche of polyploids could be obtained via higher self-pollination rates, reducing reliance on pollinators (Rodger & Ellis, 2016). Under both alternatives, by ensuring reproduction and reducing competition (Baduel *et al*., 2018), these changes in pollination niche may allow polyploids to persist in sympatry with diploids (Thompson & Lumaret, 1992; Theodoridis *et al*., 2013; Batstone *et al*., 2018; Spoelhof *et al*., 2020). Certainly, understanding the dynamics of pollination niche changes is crucial for elucidating the factors that govern the establishment and persistence of polyploid lineages with sympatry with diploids (Casazza *et al*., 2017; Segraves, 2017) and ultimately how they integrate into and affect their communities. Given the varied avenues for persistence described above, direct assessment of the impact of polyploids at the community level is needed.

A broad characterization of the role of polyploids within their communities can be obtained from plant-pollinator interaction networks (Vazquez *et al*., 2012), which provide insights into community-wide structural properties. The effect of polyploidy on network structure may be exerted directly or indirectly through its effect on associated floral traits. Using plant-pollinator networks, several floral traits, such as flower shape and size, have been associated with the pollination niche, albeit at local scale (Lázaro *et al*., 2020). Because functional traits are the main forces driving the structural properties of plant-pollinator networks (Olesen *et al*., 2007; Martín González *et al*., 2012) and polyploidy mediates a suite of floral differences and potentially niche breath, we predict that unique roles of polyploids will cascade to effect overall network structure, as illustrated in Fig. 1. The combined knowledge of plant-pollinator networks with plant ploidy classifications will facilitate understanding of how polyploid frequency influences the structure of plant-pollinator communities (Memmott *et al*., 2004; Blüthgen *et al*., 2006; Olesen *et al*., 2007; Bascompte & Stouffer, 2009; Thébault & Fontaine, 2010; Martín González *et al*., 2012). However, the multidimensional nature of phenotypes may buffer the predictive power of floral traits on the pollination niche, particularly across varying localities (Ollerton *et al*., 2009). Ultimately, to understand the effect of polyploidy on communities, the association between floral traits, polyploidy, and plant-pollinator interaction needs to be explored.

**Figure 1.**
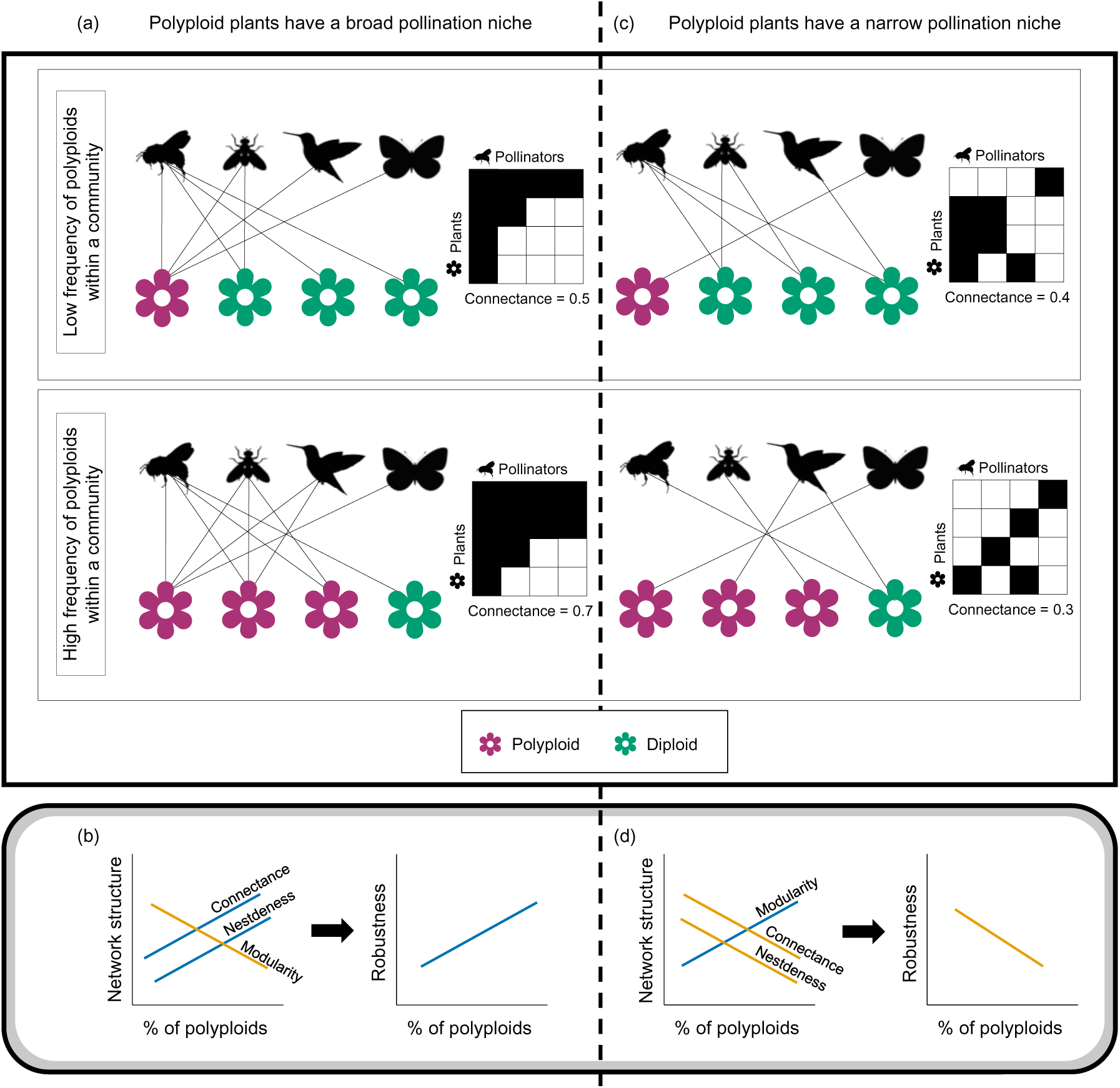
Examples of how the frequency of polyploid plants in a community is hypothesized to impact network structure under different pollination niche scenarios. In scenario (a), polyploids have a broader pollination niche relative to diploids, thus (b) the increase of polyploid frequency in a community is expected to increase the network indices connectance, nestedness, that can confer higher robustness to the community. In scenario (c), polyploids have a narrower pollination niche relative to diploids, thus (d) the increase of polyploid frequency is expected to increase modularity but reduce connectance which could cascade to a less robust community. Polyploid plants are represented as pink flowers and diploid plants as green flowers. Orange lines correspond to negative relations and blue to positive ones (b,d).

Network-wide indices, including connectance, nestedness, and modularity, capture key structural aspects of mutualistic interaction networks and are predicted to influence community robustness (Duan *et al*., 2023). A network with high connectance represents a richer ecological community, with a high degree of mutualistic interactions. Nestedness measures the extent to which specialists interact with subsets of the species that generalists interact with (Bascompte *et al*., 2003). High nestedness may indicate reduced competition, potentially enhancing the resilience of the community to disturbances (Fortuna *et al*., 2010), although the effect of nestedness depends on the nature of plant phenotypes within the community (e.g., co-flowering plants sharing a pollinator could compete with one another). Modularity quantifies the tendency of a network to be composed of tightly interconnected substructures (Newman & Girvan, 2004; Guimerà & Amaral, 2005). High modularity can enhance stability by compartmentalizing the effects of disturbances. However, it can also hinder adaptability and increase vulnerability if the modules are too isolated (Xiang et al., 2023).

Here, we provide a comprehensive assessment of the relationship between the frequency of polyploids within their communities and the structure of plant-pollinator networks. We hypothesize that if polyploids, in general, have broader pollination niches than diploids (Fig. 1a), then increased presence of polyploids is expected to enhance aspects of network structure, such as connectance, that promote stability of the community (Okuyama & Holland, 2008; Martín González *et al*., 2010; Thébault & Fontaine, 2010; Johnson & Ashman, 2019). This could also lead to increased nestedness and decreased modularity, particularly in larger networks (Fig. 1b) (Fortuna *et al*., 2010; Landi *et al*., 2018). On the other hand, a narrowed polyploid pollination niche (Fig. 1c) should result in polyploids having more peripheral roles within their communities, increased modularity, and reduced connectance and robustness in the face of extinction (Fig. 1d) (Thébault & Fontaine, 2010; Theodoridis *et al*., 2013; Baduel *et al*., 2018; Spoelhof *et al*., 2020). Furthermore, we hypothesize that the effect of polyploidy on network structure could be mediated through associated functional traits. Specifically, higher frequency of self-compatible polyploids may increase specialization of the network, that is increasing modularity and decreasing nestedness and connectance of the community. On the other hand, increased frequency of polyploids exhibiting flower shape that is permissible to pollinators would contribute to generalization aspects of the associated networks. Namely, reducing modularity and increasing nestedness and connectance.

To examine the role of polyploids within their ecological communities, we assembled an extensive dataset of 316 plant-pollinator networks, spanning 7,374 unique pollinators and 2,007 plant species of which 1,448 have ploidy classification. These were supplemented with environmental data as well as data on mating system and flower shape. In our analysis we chose to characterize flower shape as ‘restrictive’ or not because it directly reflects the potential for pollinators to access floral rewards and mediate pollination (i.e., it is a function of a variety of floral morphological traits (Burns *et al*., 2019)). Mating system (categorized here as self-compatible or incompatible) facilitates polyploid persistence and self-compatible plants were shown to have narrower pollination niche (Streher et al., 2023). We further included environmental factors known to influence polyploidy (Rice *et al*., 2019) or plant-pollinator networks (Liu *et al*., 2021). With these data we aim to answer the following questions: (1) How does the presence of polyploids affect the structure of their plant-pollinator communities? (2) Which reproductive traits and environmental factors mediate these effects? (3) Does an increased prevalence of polyploids impact the network resilience to extinction?

## Materials and methods

### Assembly of plant-pollinator networks and trait data

A plant-pollinator community can be represented by a connectivity matrix depicting the presence/absence of visits by pollinator taxa to a plant species (denoted as a binary network) or the frequency of pollinator visits (denoted as a weighted network). Here, we refer to plant-pollinator interaction networks although we acknowledge some published networks in our data set may have recorded flower visitation but not confirmed effective pollination by the visitors per se. To examine the effect of polyploidy on interactions of plants with pollinators, we collected 404 weighted networks from three online resources: Interaction Web Database (IWDB) (http://www.ecologia.ib.usp.br/iwdb/), Mangal DB (Poisot *et al*., 2016), and Web of life (Bascompte, 2009). These were supplemented with a literature search that retrieved 335 additional unique networks (Aizen *et al*., 2008; Trøjelsgaard *et al*., 2019; Schwarz *et al*., 2020). In total, 739 weighted networks were collected. The geographic locations of the plant-pollinator community from which the networks were obtained were extracted from the three online databases or, if not available, via manual data mining from the source manuscripts. We then mapped the coordinates (longitude, latitude) to climatic data in the years for 1970-2000 using the R package raster (Hijmans, 2023) and WorldClim version 2.1 (Fick & Hijmans, 2017). The WorldClim Global Climate Data consists of 19 temperature and precipitation characteristics (BIO1 to BIO19) that were developed from climate data during 1970–2000 at a ten arc-minute resolution. Four climatic factors were used in our analysis: temperature seasonality (BIO4), precipitation seasonality (BIO15), mean temperature of warmest quarter (BIO10), and precipitation of warmest quarter (BIO18). We anticipated that two of these could act as drivers of plant-pollinator interaction dynamics: BIO15 has been directly associated with increased modularity and decreased robustness (Liu *et al*., 2021); and BIO10 was shown to be strongly associated with polyploid distribution across the globe (Rice *et al*., 2019) and could consequently act as a driver of changes in plant-pollinator interactions. BIO4 and BIO18 were chosen as complementary to BIO15 and BIO10. The collected networks and the metadata assigned to them are available in supplementary files S1 and S2.

To enable integration of data from multiple sources and to assess the taxonomic resolution of the networks, name resolution was applied for both plants and pollinators appearing in the networks. Name resolution is a procedure that maps the original names recorded in an empirical dataset to their currently accepted names, while correcting for possible misspellings. Name resolution was applied using the Taxonome tool(Kluyver & Osborne, 2013) against the World Flora Online for plant names (Borsch *et al*., 2020), and against data retrieved from the Integrated Taxonomic Information System (ITIS; www.itis.gov) for pollinator names. This resulted in 79% (4,261 records) of the plant names and 21% (2,885 records) of the pollinator names resolved at the species level. Notably, most pollinator names were documented at the species level, but were resolved at genus level due to name ambiguity (e.g., *Pollenia* sp1 subsp. m_pl_006), with only 10% of the unresolved names consisting of a single word. Plant classification as either polyploid or diploid was obtained from PloiDB (Halabi *et al*., 2023) based on resolved names (or original name if unresolved). Here, we used classifications devised from inferences at the genus-level, which were found to produce the highest agreement with data reported in the literature, and mostly refer to neo- and meso-polyploids(Halabi *et al*., 2023). Additionally, to quantify the mediated effect of polyploidy through associated plant traits, we assembled a database of plant reproductive traits known to be impacted by polyploidy (mating system and flower restrictiveness (Vamosi et al., 2007; Streher et al., 2023)). Mating system data (self-compatible or self-incompatible) was collected from two main compilations (Grossenbacher *et al*., 2016; Zenil-Ferguson *et al*., 2019) and further supplemented using a literature search. Additionally, following previous studies (Burns et al., 2019; Lanuza et al., 2023),we categorized plant species into those with flower shapes that restrict pollinator access to reproductive parts or rewards (e.g., bells, tubes, spurs, flag/keel, orchid flowers, poricidal anthers, etc.) and those without such limitations (e.g., open, bowls, disk, etc.). Flower restrictiveness information was obtained primarily through species descriptions available in a variety of online sources (GBIF Occurrence Download; Borsch *et al*., 2020). When not available, we searched for information in peer-reviewed manuscripts and iNaturalist images (www.inaturalist.org; Nugent, 2018) (See additional details in Supplementary Note S1).

To mitigate sampling biases and to limit our analyses to networks of higher resolution, we included in our dataset only weighted networks that meet the following criteria: contain more than five pollinator taxa, more than five plant species with assigned ploidy information, and ploidy classifications available for at least 50% of the plant species. We additionally excluded networks for which climatic data or trait data for the plant species within them were unavailable. This filtering resulted in a dataset of 316 networks, spanning 2,007 unique plant species and 7,374 unique pollinators.

### Modeling the link of polyploid frequency with community structure

#### Measures of community structure

To test whether polyploid frequency within a community affects network architecture, we computed four network indices across the set of weighted networks collected here. These indices capture key structural aspects of the interaction network: (1) Connectance, defined as the proportion of realized interactions out of all possible interactions across species within the network. It represents the actual connections formed compared to the total potential connections in the network. To compute this measure, all networks were treated as binarized; (2) Nestedness, defined as the degree to which the network consists of nested substructures, in which specialist members primarily interact with members that generalists also interact with, and computed as weighted NODF (nested overlap and decreasing fill (Almeida-Neto *et al*., 2008)) (3) Modularity, defined as the as the tendency of a network to be organized in modules, such that members within a module mostly interact among themselves and little with members from other modules, and computed using the Q index (Newman & Girvan, 2004). The modules within each network were detected using the optimization algorithm of Beckett (2016); (4) Robustness, defined as the ability of a community to endure extinctions of its members, and computed as the area under the extinction curve under random extinction of plant species, following Burgos et al. (2007). Following previous studies (Escobedo-Kenefic *et al*., 2020; Stein *et al*., 2021; Vollstädt *et al*., 2022; Herrmann *et al*., 2023), we delta-transformed all indices relative to a null distribution (i.e., subtracted the mean across null simulated networks from the raw values). The null simulations were conducted while preserving the network size using the algorithm of Patefield (1981). The null simulations and computation of all indices were obtained using the bipartite R package (Dormann *et al*., 2009).

#### Statistical analysis

The polyploid frequency within a network (hereafter, %PP) is defined here as the proportion of plant species classified as polyploids out of all the plant species within the network for which a ploidy classification was available. We first conducted a regression analysis to examine the effect of %PP, as well as that of other reproductive traits and environmental factors, on the computed network indices. This was carried out independently for each of the four network indices (connectance, nestedness, modularity, and robustness). Specifically, we initially examined the following predictors corresponding to reproductive trait frequencies within the networks: frequency of restrictive flowers (hereafter %Restrictive) and of self-compatible plants (hereafter %SC). Similar to %PP, reproductive trait frequencies were computed relative to the number of plant species within the network for which a trait value was available. As such, each frequency value possesses a varying degree of missing data. The four environmental factors (BIO4, BIO10 – corresponding to temperature seasonality and mean value at the warmest quarter, and BIO15, BIO18 – corresponding to similar measures of precipitation) were also included. To overcome skewness, and following a previous study (Poppenwimer *et al*., 2023), the precipitation variables were log transformed.

To assess the association of each variable on each network-based index, univariate linear models were fitted to the different response indices (connectance, nestedness, modularity, robustness), each time with a different predictor. To account for spatial dependencies among samples, we tested the residuals of each model for spatial autocorrelation using the R package terra (Hijmans R, 2023). A significant positive spatial autocorrelation was detected for all models examined, and we thus fitted additional simultaneous autoregressive (SAR) models (Kissling & Carl, 2008) under the same settings using the R package spdep (Bivand Roger, 2022).

While univariate regression analysis can identify associations between individual predictor variables and network structure, it cannot determine their relative contributions or distinguish between direct and indirect effects. To uncover the relative effects of %PP on network structure, either direct or mediated via plant traits, we conducted a path analysis, using the R package piecewiseSEM (Lefcheck, 2016). Path analysis is a statistical technique used for analyzing complex relationships among variables by simultaneously examining the interdependencies between them and enabling exploration of their causal relationships. For each network index, we constructed a structural equational model with a diagram that was based on theoretical assumptions and was identical for all the examined network indices (Fig. 3). In these diagrams, %PP was assumed to have a direct effect on the examined network index, and indirect effects mediated via %SC and %Restrictive. Additionally, all plant factors (%PP, %SC and %Restrictive) and the examined network index were assumed to be affected by environmental conditions. To avoid using excessively complicated models, in this analysis we included a single environmental variable (BIO15: precipitation seasonality), which had the overall highest contribution according to multivariate regression analyses (Supplementary Note S2). Additionally, we included network size in the diagram to account for variability in network indices related to network size and accommodated for environmental influence on network size as an edge in the diagram. During fitting, we verified that no additional edges should be added to the diagram using d-separation tests (Shipley, 2013). Finally, to assess the robustness of the path analysis, we explored several variations to this initial model: (1) using BIO10 (mean temperature of warmest quarter; the most influential temperature-based variable in the multivariate analysis) as an alternative environmental factor; (2) including both BIO10 and BIO15 as environmental predictors; (3) excluding the %Restrictive factor from the diagram; (4) excluding the %SC factor from the diagram.

## Results

### Network data assembly

We assembled an extensive dataset of plant-pollinator interaction networks and supplemented these with ploidy estimates. Initially, a total of 739 weighted networks were retrieved. Following data filtration to ensure the analyses of networks of sufficient quality, our final dataset consisted of 316 networks that were globally distributed (Supplementary Fig. S1), but somewhat biased toward temperate northern localities, reflecting the uneven global distribution of studies biased towards non-tropical areas (Vizentin-Bugoni *et al*., 2018). The majority of the analyzed networks comprised of fewer than 20 plant species (average = 18) and 60 pollinators (average = 52), overall encompassing 5,659 plant vertices and 16,846 pollinator vertices, corresponding to 2,007 unique plant species, belonging to 142 families and 7,374 unique pollinator names. For plant species, we were able to obtain ploidy classifications for 1,448 species (the %PP across networks ranged from 5% to 100%), flower restrictiveness for 1,148 species, and mating system for 543 species. The distributions of the plant traits across the analyzed networks are shown in Fig. S2, and the trait information per species is available in Supplementary File S3.

### The association of polyploid frequency with community structure

To quantify the effect of polyploid frequency as well as other potential factors on network structure, we initially conducted a univariate regression analysis. This was carried out independently for four network indices that characterize different aspects of network structure: connectance, nestedness, modularity, and robustness. These analyses showed that frequency of polyploids within networks (%PP) was associated positively with nestedness (*p* = 0.004), and negatively with modularity (*p*= 0.006), while the associations with connectance and robustness were positive albeit statistically non-significant (Fig. 2; Table 1). The non-significant impact of polyploids on robustness was further supported by additional extinction simulations (varying which plants went extinct first), which indicated that the robustness of a network remains largely unaffected by the ploidy of plants selected for primary extinction (Supplementary Note S4). Among the related factors examined, the frequency of self-compatible plants (%SC) was the most influential – being significantly associated with all four indices examined (*p* < 0.001). The predictive power of the percentage of restrictive flowers (%Restrictive) was generally less prominent and significant only for modularity (*p* = 0.003). Interestingly, in all four indices the associations of %SC and %Restrictive were in the opposite direction to that observed for %PP. Among the environmental factors examined, networks with lower connectance and robustness were generally located in regions with warm summers (i.e., higher values of BIO10; *p* = 0.049 and 0.299 for connectance and robustness, respectively) and dry summers (i.e., higher values of BIO18; *p* = 0.016 and 0.011 for connectance and robustness, respectively). Moreover, lower connectance and robustness were observed in regions with high seasonality (BIO4 and BIO15: *p* < 10^−4^). While environmental associations with nestedness and modularity were generally less prominent, networks in regions of higher precipitation seasonality (BIO15) had significantly lower levels of nestedness (Table 1). The direction of all associations remained unchanged upon replacing linear models with spatial autoregressive models (Table S1), although some of the associations became non-significant, suggesting that the spatial effect may be stronger than the predictive power of some examined factors. The patterns depicted above were mostly corroborated using a multivariate regression analysis. As may be expected, the associations of %PP somewhat weakened when considering other factors within the same statistical framework, although its effect on network nestedness decreased only slightly and remained significant (𝑝 = 0.011, Supplementary Table S2; Supplementary Notes S2).

**Figure 2.**
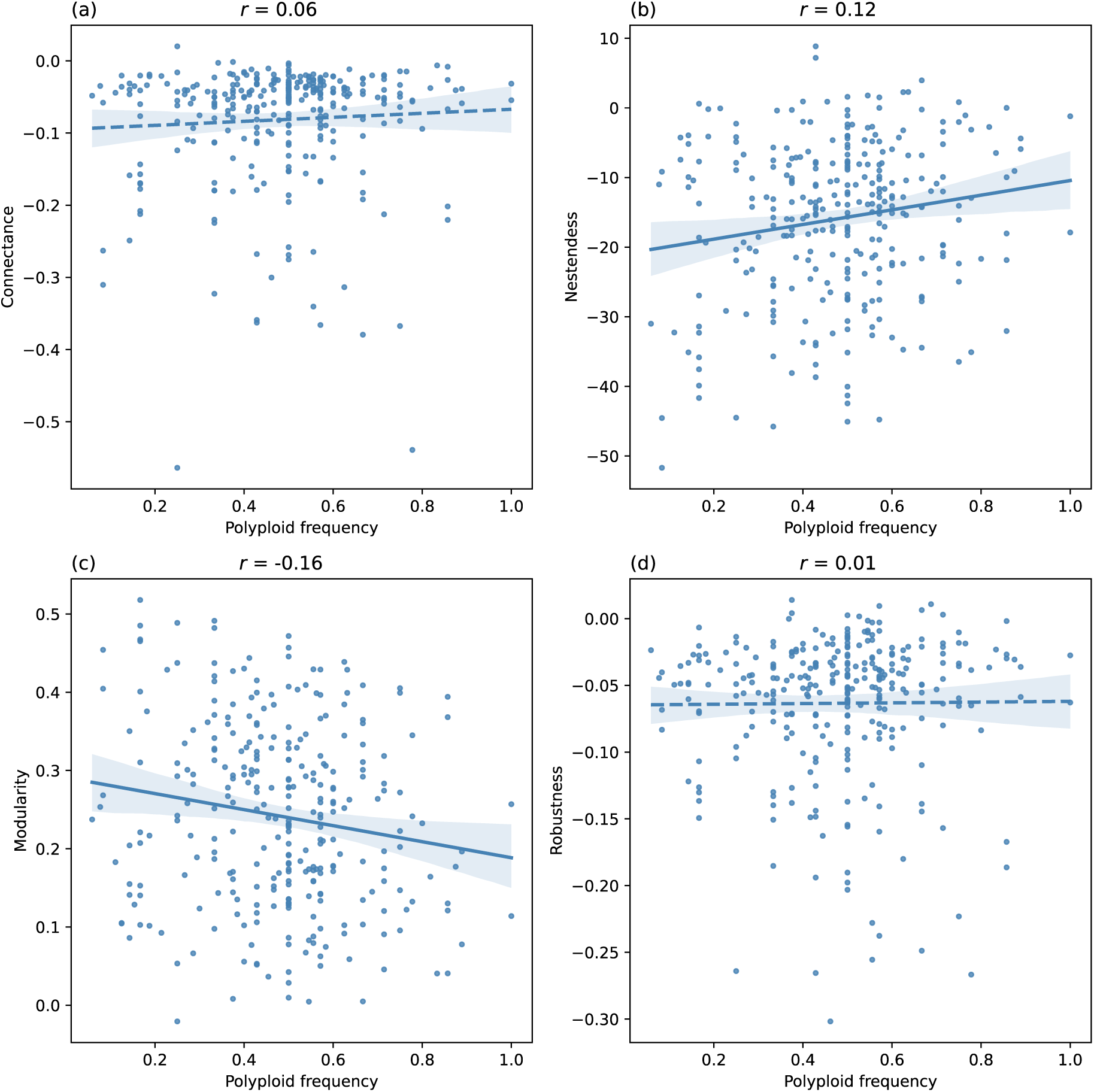
The distribution of standardized plant-pollinator network metrics plotted across with increasing frequencies of polyploid taxa. Metrics of connectance (a), nestedness (b), modularity (c), and robustness (d) are shown across the 316 weighted networks included in the analysis. The Pearson correlation coefficient, r, is listed above each plot.

**Table 1.** Results of the univariate regression analysis. . For each network-based index, the Pearson correlation coefficient (𝑹) and its standardized value (𝜷) are displayed, along with the 𝒑-value of the respective regression model. Significant associations are bolded.

| Predictor | Connectance |  | Nestedness |  | Modularity |  | Robustness |  |
| --- | --- | --- | --- | --- | --- | --- | --- | --- |
|  | *R* (*β*) | *p* | *R* (*β*) | *p* | *R* (*β*) | *p* | *R* (*β*) | *p* |
| %PP | 0.06 (1.02) | 0.306 | **0.17 (2.91)** | **0.004** | **-0.16 (-2.75)** | **0.006** | 0.009 (0.16) | 0.875 |
| %SC | **-0.19 (-3.38)** | **0.001** | **-0.18 (-3.09)** | **0.002** | **0.17 (3.04)** | **0.002** | **-0.17 (-2.93)** | **0.004** |
| %Restrictive | -0.05 (-0.9) | 0.366 | -0.01 (-0.18) | 0.852 | **0.17 (2.95)** | **0.003** | -0.01 (-0.17) | 0.862 |
| BIO4<sup>a</sup> | **-0.32 (-5.83)** | **<10<sup>-4</sup>** | -0.09 (-1.62) | 0.104 | 0.06 (1.02) | 0.306 | **-0.30 (-5.51)** | **<10<sup>-4</sup>** |
| BIO10<sup>b</sup> | **-0.11 (-1.96)** | **0.049** | -0.04 (-0.61) | 0.536 | 0.10 (1.75) | 0.08 | -0.06 (-1.04) | 0.299 |
| BIO15<sup>c</sup> | **-0.33 (-6.07)** | **<10<sup>-4</sup>** | **-0.15 (-2.67)** | **0.008** | 0.03 (0.53) | 0.59 | **-0.32 (-5.82)** | **<10<sup>-4</sup>** |
| BIO18<sup>d</sup> | **0.14 (2.4)** | **0.016** | 0.02 (0.34) | 0.732 | -0.01 (-0.23) | 0.82 | **0.17 (2.54)** | **0.011** |
| Network size | **0.30 (5.4)** | **<10<sup>-4</sup>** | **0.12 (2.15)** | **0.032** | **0.27 (4.89)** | **<10<sup>-4</sup>** | **0.19 (3.41)** | **0.001** |

<sup>a</sup>Temperture seasonality; <sup>b</sup>Mean temperature of warmest quarter; <sup>c</sup>Precipitation seasonality; <sup>d</sup>Precipitation of warmest quarter.

### The mediated effect of polyploid frequency on community structure

The results of the path analysis were consistent with those derived from the univariate regression analysis, but also revealed the interplay between direct and indirect effects of key factors affecting network structure. As shown in figure 3, the path analysis revealed several key findings. First, positive direct effects of %PP on connectance, nestedness, and robustness, and a negative effect on modularity were observed. Second, among the trait-based attributes, %SC had the strongest impact on network structure, such that higher %SC led to lower connectance, nestedness, and robustness, but higher modularity. The effect of %Restrictive was weaker than that of %SC, albeit in the case of modularity, it had a significant positive direct effect. This, coupled with the effect of %SC, had a combined significant contribution to the increase of network compartmentalization. Third, higher %PP was negatively associated with both %SC and %Restrictive. Moreover, across all network indices examined, the direct effect of %PP was in the opposite direction to that of %SC and %Restrictive. Fourth, in the case of nestedness, the effect of %PP was significant and stronger in magnitude than that of %SC. This was not the case for connectance, modularity, and robustness, where the effect of %PP was weak and substantially lower (in absolute terms) than that of %SC. Fifth, decomposing the total effect of %PP into the direct and indirect effects (Table 2), revealed the multiple ways %PP affected network structure. For example, while the direct effect of %PP on nestedness was much larger than its indirect effects via both %SC and %Restrictive (direct effect of 0.174 versus combined indirect effect of 0.02 [-0.005+0.027]), this was not the case for the other indices where the combined indirect effect of %PP was stronger (four times so for robustness) or similar (connectance and modularity) to the direct effects. Finally, the analysis indicated that climate had a prominent effect on network size, with a negative association with precipitation seasonality (BIO15), indicating that lower precipitation seasonality is linked with higher species richness. Additionally, precipitation seasonality had a strong direct negative effect on network connectance and robustness. Similar overall results were obtained when the effect of temperature (BIO10) was examined instead or in addition to precipitation seasonality (Fig. S3 and Fig. S4). In addition, the direction and magnitude of the environmental effects as well as those of %PP were similar when using simpler models, from which either the %SC or %Restrictive were excluded (Fig. S5). Further details on all model variants are given in Supplementary Note S3.

**Table 2.** Direct, indirect and total effects of polyploid frequency (%PP) on the four examined plant-pollinator network indices. The total effects of indirect paths via frequency of self-compatible (%SC) and frequency of restrictive flowers (%Restrictive) computed in the path analysis (Fig. 3) are shown.

| Response | Direct / indirect | Effect<sup>a</sup> |
| --- | --- | --- |
| Connectance | Direct effect of %PP | 0.061 |
|  | Indirect via %SC | 0.035 |
|  | Indirect via %Restrictive | 0.027 |
|  | Total effect of %PP | 0.123 |
| Nestedness | Direct effect of %PP | 0.174 |
|  | Indirect via %SC | 0.027 |
|  | Indirect via %Restrictive | -0.005 |
|  | Total effect of %PP | 0.196 |
| Modularity | Direct effect of %PP | -0.063 |
|  | Indirect via %SC | -0.033 |
|  | Indirect via %Restrictive | -0.039 |
|  | Total effect of %PP | -0.135 |
| Robustness | Direct effect of %PP | 0.011 |
|  | Indirect via %SC | 0.033 |
|  | Indirect via %Restrictive | 0.015 |
|  | Total effect of %PP | 0.059 |

<sup>a</sup>Direct paths are the standardized coefficients ( $\beta$ ); indirect paths are the product of the standardized coefficients across their respective edges; total effect is the sum of all effects.

## Discussion

Our analysis of hundreds of plant-pollinator networks demonstrated that polyploidy, a profound and widespread evolutionary process in plants, can impact key aspects of network structure. These impacts were at times subtle, and the path analysis revealed that these are often mediated by reproductive traits rather than through the direct effect of polyploidy per se. Our analysis thus uncovered the nuanced and complex way evolution can impact biotic communities. Specifically, we found that networks with higher polyploid frequency have more nested and less modular structures. These effects, however, do not appear to propagate into increased connectance and enhanced robustness to extinction. Our path analyses further revealed that the impacts of polyploid frequency are often mediated through the frequency of self-compatible plants and restrictiveness of flowers within the networks, and that these contribute to significant increase in nestedness and decrease in modularity. Moreover, the climatic effects on network structure revealed here, with precipitation seasonality being associated negatively with connectance, nestedness and robustness, and positively with modularity, confirms previous observations obtained at a smaller scale for robustness and modularity (Liu *et al*., 2021). The analysis further revealed a direct positive effect of precipitation seasonality on polyploid frequency, suggesting that animal-pollinated plant communities in regions with higher oscillations in precipitation tend to be more polyploid-rich.

The increased nestedness and decreased modularity in polyploid-rich networks suggest that polyploids experience an expansion of the pollination niche. Specifically, the results indicated that high polyploid frequency within networks confers more hierarchical substructures, potentially those in which polyploids interact with both generalist and specialist pollinators, rather than modular substructures in which plants interact with a specific subset of pollinators (Fig. 1). This coupling of decreased modularity and increased nestedness was previously shown to be prevalent in large and highly connected networks (Fortuna *et al*., 2010), our analysis adds a mechanistic dimension to this finding as we controlled for network size. While increased nestedness has been linked with increased robustness to species extinction within mutualistic networks (Duan *et al*., 2023), our results indicated that polyploid frequency has little effect of robustness. Possibly, the influence of polyploidy within the studied networks may have been moderated through factors not accommodated for in our study (Gómez *et al*., 2023). As such, multiple opposing effects on network resilience might obscure one another, resulting in an overall weak and negligible association of polyploidy with robustness, despite the observed positive association with nestedness.

Our results suggest that higher polyploid frequency is associated with changes in trait frequencies that affect community structure. Our results suggest that higher polyploid frequency is associated with decreased frequencies of self-compatible plants and of those with restrictive flowers, both being traits influential to community structure, and in particular lower frequency of these types of species in the community promote networks of less compartmentalized structures (Fig 3). Self-compatibility, in particular, has been hypothesized to be common among polyploid plants (Stebbins, 1950), but this hypothesis has not been supported at large scales (Mable, 2004) nor in a community context. We acknowledge, however, that other floral traits have been associated with pollinator accessibility (Stebbins, 1970; Bradshaw & Schemske, 2003; Faegri & L. Van der Pijl., 2013; Cortés-Flores *et al*., 2017; Lázaro *et al*., 2020), and such traits may also be associated with polyploidy. For example, flower size has been hypothesized to be larger in polyploids, attributed to the *gigas* effect (Ramsey & Schemske, 2003), and this trait has also been linked with a wider pollination niche based on the analysis of plant-pollinator interaction networks (Lázaro et al., 2020; Xiang et al., 2023). To date, most studies on the association between floral traits and the pollination niche have been conducted at a local scale, focusing on a specific set of plant species or a limited set of networks, with little support at a broader scale (Ollerton *et al*., 2009). Thus, while the link between polyploidy, floral traits, and pollination niche remains virtually unexplored at the species level (but see Streher *et al*., 2023), our work at the network level suggests that polyploids have different niches than diploids and this cascades to alter structural characteristics of the plant-pollinator network.

**Figure 3:**
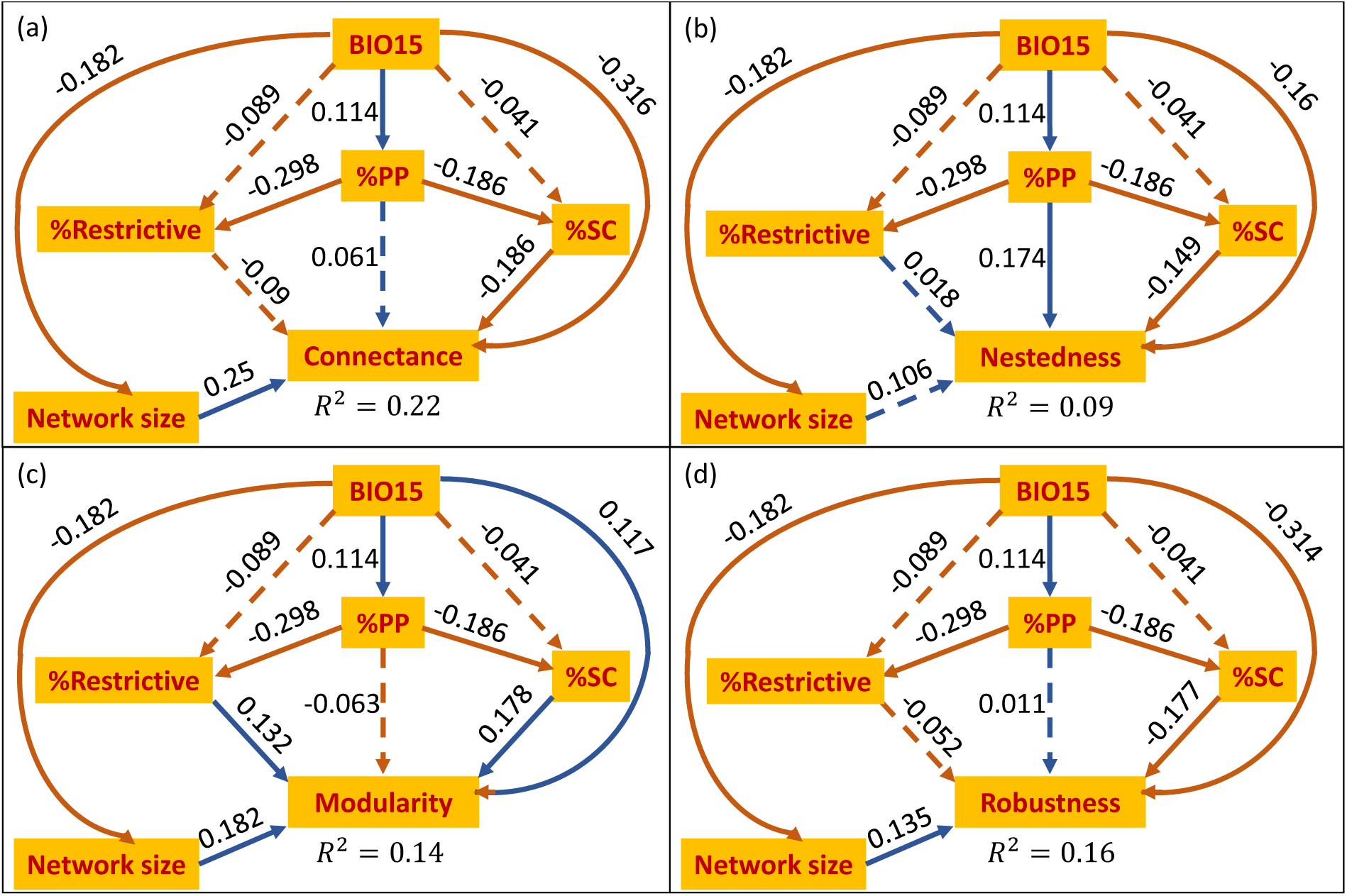
Path diagrams of the four examined network metrics connectance (a), nestedness (b), modularity (c) and robustness (d). BIO15 is precipitation seasonality, %PP corresponds to the frequency of polyploids within each network, and %SC and %Restrictive correspond to the frequencies of self-compatible plants and those with respective flower shape, respectively. Full lines correspond to paths with significant contribution and dashed lines correspond to paths with a non-significant contribution. Orange lines correspond to negative coefficients and blue to positive ones. The standardized coefficients are shown next to each edge in the diagram. The 𝑹^𝟐^is shown next to each network index. The 𝑹^𝟐^is shown next to each network index. In all panels, the results of the 𝝌^𝟐^ test for model adequacy was non-significant (p > 0.2), indicating that the model is adequate.

The analysis conducted here was performed at the network level, testing whether higher frequency of polyploids alters structural characteristics of their networks. An alternative to the network-level approach, would be to focus on the species level, in which each species within a network acts as a data point, thus potentially providing a more explicit examination on the role of polyploids within their communities. However, a meta-analysis of this type across many networks suffers from several shortcomings that have greater impact at a species-level, as compared to a network-level analysis. For example, the lack of a standardized approach in empirically recording plant-pollinator interactions introduces inherent biases, such as those attributed to sampling methods (e.g., visit-based versus pollen-based) (De Manincor *et al*., 2020), or the taxonomic resolution of the identified taxa. Such biases could significantly contribute to technical variance of the same taxon across different networks. In our analysis, we observed inconsistencies in the taxonomic resolution of the pollinators across the analyzed networks, and these inconsistencies affect the characteristics of the plant species within the networks (e.g., a plant species whose pollinators were documented at the genus level in one network may appear more specialized than in another where in which pollinators were documented at the species-level). In addition, temporal effects, which often cannot be accommodated for due to lack of information on the time in which a network was collected, may lead to variability between networks collected from the same location over time (Burkle & Alarcón, 2011; Chacoff *et al*., 2018). While such biases also impact network-level analyses, their effect is more pronounced in a species-level analysis, where each network contributes a varying number of data points. Indeed, network-level indices were shown to be less sensitive to the sampling method compared to species-level ones (Gibson *et al*., 2011). A solution to approach species-level questions of pollination niche would be focused studies of given taxonomic groups (e.g., Streher et al 2023). Such studies complement the network-level approach taken here but cannot address the community consequences of polyploidy.

Overall, our results indicate that variation in plant-pollinator community structure can be partially attributed to the relative abundance of polyploids within them. Specifically, we find that higher polyploid frequency within communities is associated with higher nestedness and lower modularity. The finding that these dependencies are mediated through floral or reproductive traits provides a much-needed mechanism for understanding how communities are generally structured. Our results demonstrate how a prominent genomic evolutionary process, such as polyploidy, plays a role in shaping ecological community structure, and highlights shifts in mating system and floral shape as important pathways through which polyploidy may impact pollinator communities in the future. Our work highlights the necessity for future research on polyploidy-driven shifts at both the community and the species levels. This will help to further explore the mechanisms behind these phenomena and understand how they might be influenced by global changes in pollinator availability and climate-driven range shifts that alter plant community composition. One might predict that if polyploids indeed exhibit greater resilience to abiotic stresses induced by climate change (van de Peer *et al*., 2021; Tossi *et al*., 2022), then plant communities will become richer with polyploids and exhibit increased nestedness and reduced modularity. Whether polyploid-rich networks will ultimately prove to be more resilient still remains to be determined.

## Acknowledgment

We thank Niv DeMalach, Anna Rice, Tyler Poppenwimmer, and Daniel B. Stouffer for their valuable suggestions and Trezalka Budinsky and Ethan Richardson for trait scoring. This study was supported by fellowships from the Tel Aviv Data Science Center (to KH) and the Edmond J. Safra Center for Bioinformatics at Tel-Aviv University (to KH and NE). This work was funded by the National Science Foundation (DEB-2027604 to TLA), the US-Israel Binational Science Foundation (BSF 2020625 to IM), and by the Israel Science Foundation (1843/21 to IM).

## Author contributions

IM and TA conceived the study. KH and NE compiled the networks dataset and wrote the code for data pre-processing and statistical analyses. NS empirically collected and compiled the plant phenotypes dataset provided consult throughout the study. TP provided consult throughout the study. IM and TA supervised the study.

## Data availability

The code for data compilation and statistical analyses is available at https://github.com/halabikeren/plant_pollinator_networks/. The compiled data is available upon request from the authors.

## References

Aizen MA, Morales CL, Morales JM. 2008. Invasive Mutualists Erode Native Pollination Webs. PLOS Biology 6: e31.

Almeida-Neto M, Guimarães P, Guimarães PR, Loyola RD, Ulrich W. 2008. A consistent metric for nestedness analysis in ecological systems: reconciling concept and measurement. Oikos 117: 1227–1239.

Anneberg TJ, Turcotte MM, Ashman TL. 2023. Plant neopolyploidy and genetic background differentiate the microbiome of duckweed across a variety of natural freshwater sources. Molecular Ecology 32: 5849–5863.

Ashman TL, Knight TM, Steets JA, Amarasekare P, Burd M, Campbell DR, Dudash MR, Johnston MO, Mazer SJ, Mitchell RJ, et al. 2004. Pollen limitation of plant reproduction: Ecological and evolutionary causes and consequences. Ecology 85: 2408–2421.

Baduel P, Bray S, Vallejo-Marin M, Kolář F, Yant L. 2018. The ‘Polyploid Hop’: Shifting challenges and opportunities over the evolutionary lifespan of genome duplications. Frontiers in Ecology and Evolution 6.

Baniaga AE, Marx HE, Arrigo N, Barker MS. 2020. Polyploid plants have faster rates of multivariate niche differentiation than their diploid relatives. Ecology Letters 23: 68–78.

Bascompte J. 2009. Disentangling the web of life. Science 325: 416–419.

Bascompte J, Jordano P, Melián CJ, Olesen JM. 2003. The nested assembly of plant-animal mutualistic networks. PNAS 100: 9383–9387.

Bascompte J, Scheffer M. 2022. The Resilience of Plant-Pollinator Networks. Annu. Rev. Entomol. 2023 68: 363–380.

Bascompte J, Stouffer DB. 2009. The assembly and disassembly of ecological networks. Philosophical Transactions of the Royal Society B: Biological Sciences 364: 1781–1787.

Batstone RT, Carscadden KA, Afkhami ME, Frederickson ME. 2018. Using niche breadth theory to explain generalization in mutualisms. Ecology 99: 1039–1050.

Beckett SJ. 2016. Improved community detection in weighted bipartite networks. Royal Society Open Science 3.

Bergamo PJ, Streher NS, Wolowski M, Sazima M. 2020. Pollinator-mediated facilitation is associated with floral abundance, trait similarity and enhanced community-level fitness. Journal of Ecology 108: 1334–1346.

Bivand Roger. 2022. R Packages for Analyzing Spatial Data: A Comparative Case Study with Areal Data.”. 54: 488–518.

B. Lanuza J, Rader R, Stavert J, Kendall LK, Saunders ME, Bartomeus I. 2023. Covariation among reproductive traits in flowering plants shapes their interactions with pollinators. Functional Ecology 37: 2072–2084.

Blüthgen N, Menzel F, Blüthgen N. 2006. Measuring specialization in species interaction networks. BMC Ecology 6: 1–12.

Borsch T, Berendsohn W, Dalcin E, Delmas M, Demissew S, Elliott A, Fritsch P, Fuchs A, Geltman D, Güner A, et al. 2020. World Flora Online: Placing taxonomists at the heart of a definitive and comprehensive global resource on the world’s plants. Taxon 69: 1311–1341.

Bradshaw HD, Schemske DW. 2003. Allele substitution at a flower colour locus produces a pollinator shift in monkeyflowers. Nature 2003 426:6963 426: 176–178.

Brittingham HA, Koski MH, Ashman TL. 2018. Higher ploidy is associated with reduced range breadth in the Potentilleae tribe. American Journal of Botany 105: 700–710.

Burgos E, Ceva H, Perazzo RPJ, Devoto M, Medan D, Zimmermann M, María Delbue A. 2007. Why nestedness in mutualistic networks? Journal of Theoretical Biology 249: 307–313.

Burkle LA, Alarcón R. 2011. The future of plant-pollinator diversity: Understanding interaction networks across time, space, and global change. American Journal of Botany 98: 528–538.

Burns JH, Bennett JM, Li J, Xia J, Arceo-Gómez G, Burd M, Burkle LA, Durka W, Ellis AG, Freitas L, et al. 2019. Plant traits moderate pollen limitation of introduced and native plants: a phylogenetic meta-analysis of global scale. New Phytologist 223: 2063–2075.

Casazza G, Boucher FC, Minuto L, Randin CF, Conti E. 2017. Do floral and niche shifts favour the establishment and persistence of newly arisen polyploids? A case study in an Alpine primrose. Annals of Botany 119: 81–93.

Chacoff NP, Resasco J, Vázquez DP. 2018. Interaction frequency, network position, and the temporal persistence of interactions in a plant–pollinator network. Ecology 99: 21–28.

Cortés-Flores J, Hernández-Esquivel K, González-Rodríguez A, Ibarra-Manríquez G. 2017. Flowering phenology, growth forms, and pollination syndromes in tropical dry forest species: Influence of phylogeny and abiotic factors. American Journal of Botany 104: 39–49.

Cui L, Wall PK, Leebens-Mack JH, Lindsay BG, Soltis DE, Doyle JJ, Soltis PS, Carlson JE, Arumuganathan K, Barakat A, et al. 2006. Widespread genome duplications throughout the history of flowering plants. Genome research 16: 738–749.

Doré M, Fontaine C, Thébault E. 2021. Relative effects of anthropogenic pressures, climate, and sampling design on the structure of pollination networks at the global scale. Global Change Biology 27: 1266–1280.

Dormann CF, Fründ J, Blüthgen N, Gruber B. 2009. Indices, Graphs and Null Models: Analyzing Bipartite Ecological Networks. The Open Ecology Journal 2: 7–24.

Duan D, Zhai Y, Hou G, Zhou M, Rong Y. 2023. Effect of network nestedness on stability, diversity, and resilience of ecosystems. Chaos 33.

Escobedo-Kenefic N, Landaverde-González P, Theodorou P, Cardona E, Dardón MJ, Martínez O, Domínguez CA. 2020. Disentangling the effects of local resources, landscape heterogeneity and climatic seasonality on bee diversity and plant-pollinator networks in tropical highlands. Oecologia 194: 333–344.

Faegri K, L. Van der Pijl. 2013. Principles of Pollination Ecology. Elsevier.

Fick SE, Hijmans RJ. 2017. WorldClim 2: new 1-km spatial resolution climate surfaces for global land areas. International Journal of Climatology 37: 4302–4315.

Fortuna MA, Stouffer DB, Olesen JM, Jordano P, Mouillot D, Krasnov BR, Poulin R, Bascompte J. 2010. Nestedness versus modularity in ecological networks: two sides of the same coin? Journal of Animal Ecology 79: 811–817.

Gaynor ML, Ng J, Laport RG. 2018. Phylogenetic structure of plant communities: Are polyploids distantly related to co-occurring diploids? Frontiers in Ecology and Evolution 6: 52.

GBIF Occurrence Download. (GBIF, 2021).

Gibson RH, Knott B, Eberlein T, Memmott J. 2011. Sampling method influences the structure of plant-pollinator networks. Oikos 120: 822–831.

Gómez JM, González-Megías A, Armas C, Narbona E, Navarro L, Perfectti F. 2023. The role of phenotypic plasticity in shaping ecological networks. Ecology Letters 26: S47–S61.

Grossenbacher D, Briscoe Runquist RD, Goldberg EE, Brandvain Y. 2016. No association between plant mating system and geographic range overlap. American Journal of Botany 103: 110–117.

Guimerà R, Amaral LAN. 2005. Functional cartography of complex metabolic networks. Nature 2005 433:7028 433: 895–900.

Halabi K, Shafir A, Mayrose I. 2023. PloiDB: The plant ploidy database. New Phytologist 240: 918–927.

Herrmann J, Buchholz S, Theodorou P. 2023. The degree of urbanisation reduces wild bee and butterfly diversity and alters the patterns of flower-visitation in urban dry grasslands. Scientific Reports 13.

Hijmans RJ. 2023. Geographic Data Analysis and Modeling. R package raster version 3.6-26.

Hijmans R. 2023. terra: Spatial Data Analysis. R package version 1.7-46.

Integrated Taxonomic Information System (ITIS). 2019.

Jiao Y, Wickett NJ, Ayyampalayam S, Chanderbali AS, Landherr L, Ralph PE, Tomsho LP, Hu Y, Liang H, Soltis PS, et al. 2011. Ancestral polyploidy in seed plants and angiosperms. Nature 2011 473:7345 473: 97–100.

Johnson AL, Ashman TL. 2019. Consequences of invasion for pollen transfer and pollination revealed in a tropical island ecosystem. New Phytologist 221: 142–154.

Johnson CA, Dutt P, Levine JM. 2022. Competition for pollinators destabilizes plant coexistence. Nature 607: 721–725.

Kissling WD, Carl G. 2008. Spatial autocorrelation and the selection of simultaneous autoregressive models. Global Ecology and Biogeography 17: 59–71.

Kluyver TA, Osborne CP. 2013. Taxonome: a software package for linking biological species data. Ecology and Evolution 3: 1262–1265.

Köhler C, Mittelsten Scheid O, Erilova A. 2010. The impact of the triploid block on the origin and evolution of polyploid plants. Trends in Genetics 26: 142–148.

Landi P, Minoarivelo HO, Brännström Å, Hui C, Dieckmann U. 2018. Complexity and stability of ecological networks: a review of the theory. Population Ecology 60: 319–345.

Lázaro A, Gómez-Martínez C, Alomar D, González-Estévez MA, Traveset A. 2020. Linking species-level network metrics to flower traits and plant fitness. Journal of Ecology 108: 1287–1298.

Lefcheck JS. 2016. piecewiseSEM: Piecewise structural equation modelling in r for ecology, evolution, and systematics. Methods in Ecology and Evolution 7: 573–579.

Levin DA. 1983. Polyploidy and Novelty in Flowering Plants. The American Naturalist 122: 1– 25.

Liu H, Liu Z, Zhang M, Bascompte J, He F, Chu C. 2021. Geographic variation in the robustness of pollination networks is mediated by modularity. Global Ecology and Biogeography 30: 1447–1460.

Lundgren R, Totland Ø, Lázaro A. 2016. Experimental simulation of pollinator decline causes community-wide reductions in seedling diversity and abundance. Ecology 97: 1420–1430.

Mable BK. 2004. Polyploidy and self-compatibility: Is there an association? New Phytologist 162: 803–811.

De Manincor N, Hautekèete N, Mazoyer C, Moreau P, Piquot Y, Schatz B, Schmitt E, Zélazny M, Massol F. 2020. How biased is our perception of plant-pollinator networks? A comparison of visit- and pollen-based representations of the same networks. Acta Oecologica 105: 103551.

Martín González AM, Allesina S, Rodrigo A, Bosch J. 2012. Drivers of compartmentalization in a Mediterranean pollination network. Oikos 121: 2001–2013.

Martín González AM, Dalsgaard B, Olesen JM. 2010. Centrality measures and the importance of generalist species in pollination networks. Ecological Complexity 7: 36–43.

Martin SL, Husband BC. 2009. Influence of phylogeny and ploidy on species ranges of North American angiosperms. Journal of Ecology 97: 913–922.

Memmott J, Waser NM, Price M V. 2004. Tolerance of pollination networks to species extinctions. Proceedings of the Royal Society B: Biological Sciences 271: 2605–2611.

Mustajärvi K, Siikamäki P, Rytkönen S, Lammi A. 2001. Consequences of Plant Population Size and Density for Plant-Pollinator Interactions and Plant Performance. Journal of Ecology 89: 80–87.

Newman MEJ, Girvan M. 2004. Finding and evaluating community structure in networks. Physical Review E 69: 026113.

Okuyama T, Holland JN. 2008. Network structural properties mediate the stability of mutualistic communities. Ecology Letters 11: 208–216.

Olesen JM, Bascompte J, Dupont YL, Jordano P. 2007. The modularity of pollination networks. Proceedings of the National Academy of Sciences of the United States of America 104: 19891–19896.

Ollerton J, Alarcon R, Waser NM, Price M V., Watts S, Cranmer L, Hingston A, Peter CI, Rotenberry J. 2009. A global test of the pollination syndrome hypothesis. Annals of Botany 103: 1471–1480.

Ollerton J, Winfree R, Tarrant S. 2011. How many flowering plants are pollinated by animals? Oikos 120: 321–326.

Otto SP. 2007. The Evolutionary Consequences of Polyploidy. Cell 131: 452–462.

Padilla-García N, Šrámková G, Záveská E, Šlenker M, Clo J, Zeisek V, Lučanová M, Rurane I, Kolář F, Marhold K. 2023. The importance of considering the evolutionary history of polyploids when assessing climatic niche evolution. Journal of Biogeography 50: 86–100.

Patefield WM. 1981. Algorithm AS 159: An Efficient Method of Generating Random R × C Tables with Given Row and Column Totals.

van de Peer Y, Ashman TL, Soltis PS, Soltis DE. 2021. Polyploidy: an evolutionary and ecological force in stressful times. Plant Cell 33: 11–26.

Van De Peer Y, Mizrachi E, Marchal K. 2017. The evolutionary significance of polyploidy. Nature Reviews Genetics 18: 411–424.

Poisot T, Baiser B, Dunne JA, Kéfi S, Massol F, Mouquet N, Romanuk TN, Stouffer DB, Wood SA, Gravel D. 2016. mangal – making ecological network analysis simple. Ecography 39: 384–390.

Poppenwimer T, Mayrose I, Demalach N. 2023. Revising the global biogeography of annual and perennial plants. Nature 624: 109–114.

Ramsey J, Ramsey TS. 2014. Ecological studies of polyploidy in the 100 years following its discovery. Philosophical Transactions of the Royal Society B: Biological Sciences 369.

Ramsey J, Schemske DW. 2003. Neopolyploidy in Flowering Plants. Annual Review of Ecology and Systematics 33: 589–639.

Ranney TG. 2006. Polyploidy: From Evolution to New Plant Development Polyploidy: From Evolution to New Plant Development. *Combined proceedings international plant propagators’* society 56: 137–142.

Rice A, Šmarda P, Novosolov M, Drori M, Glick L, Sabath N, Meiri S, Belmaker J, Mayrose I. 2019. The global biogeography of polyploid plants. Nature Ecology & Evolution 2019 3:2 3: 265–273.

Rodger JG, Bennett JM, Razanajatovo M, Knight TM, Van Kleunen M, Ashman T-L, Steets JA, Hui C, Arceo-Gómez G, Burd M, et al. 2021a. Widespread vulnerability of flowering plant seed production to pollinator declines. Sci. Adv 7: 3524–3537.

Rodger JG, Bennett JM, Razanajatovo M, Knight TM, van Kleunen M, Ashman TL, Steets JA, Hui C, Arceo-Gómez G, Burd M, et al. 2021b. Widespread vulnerability of flowering plant seed production to pollinator declines. Science Advances 7: 3524–3537.

Rodger JG, Ellis AG. 2016. Distinct effects of pollinator dependence and self-incompatibility on pollen limitation in South African biodiversity hotspots. Biology Letters 12: 20160253.

Sargent RD, Ackerly DD. 2008. Plant-pollinator interactions and the assembly of plant communities. Trends in Ecology and Evolution 23: 123–130.

Schwarz B, Vázquez DP, CaraDonna PJ, Knight TM, Benadi G, Dormann CF, Gauzens B, Motivans E, Resasco J, Blüthgen N, et al. 2020. Temporal scale-dependence of plant– pollinator networks. Oikos 129: 1289–1302.

Segar ST, Fayle TM, Srivastava DS, Lewinsohn TM, Lewis OT, Novotny V, Kitching RL, Maunsell SC. 2020. The Role of Evolution in Shaping Ecological Networks. Trends in Ecology & Evolution 35: 454–466.

Segraves KA. 2017. The effects of genome duplications in a community context. New Phytologist 215: 57–69.

Segraves KA, Anneberg TJ. 2016. Species interactions and plant polyploidy. American Journal of Botany 103: 1326–1335.

Shipley B. 2013. The AIC model selection method applied to path analytic models compared using a d-separation test. Ecology 94: 560–564.

Soltis DE, Albert VA, Leebens-Mack J, Bell CD, Paterson AH, Zheng C, Sankoff D, DePamphilis CW, Wall PK, Soltis PS. 2009. Polyploidy and angiosperm diversification. American Journal of Botany 96: 336–348.

Soltis PS, Soltis DE. 2000. The role of genetic and genomic attributes in the success of polyploids. Proceedings of the National Academy of Sciences 97: 7051–7057.

Spoelhof JP, Keeffe R, McDaniel SF. 2020. Does reproductive assurance explain the incidence of polyploidy in plants and animals? New Phytologist 227: 14–21.

Stebbins GL. 1950. Variation and Evolution in Plants. Columbia Univ. Press.

Stebbins GL. 1970. Adaptive Radiation of Reproductive Characteristics in Angiosperms, I: Pollination Mechanisms. Annual review of ecology and systematics 1: 307–326.

Stein K, Coulibaly D, Balima LH, Goetze D, Linsenmair KE, Porembski S, Stenchly K, Theodorou P. 2021. Plant-pollinator networks in savannas of Burkina Faso, West Africa. Diversity 13: 1–14.

Streher NS, Budinsky T, Halabi K, Mayrose I, Ashman T-L. 2023a. The effect of polyploidy and mating system on floral size and the pollination niche in Brassicaceae. International Journal of Plant Sciences 185.

Streher NS, Budinsky T, Halabi K, Mayrose I, Ashman T-L. 2023b. The effect of polyploidy and mating system on floral size and the pollination niche in Brassicaceae. International Journal of Plant Sciences.

Thébault E, Fontaine C. 2010. Stability of ecological communities and the architecture of mutualistic and trophic networks. Science 329: 853–856.

Theodoridis S, Randin C, Broennimann O, Patsiou T, Conti E. 2013. Divergent and narrower climatic niches characterize polyploid species of European primroses in Primula sect. Aleuritia. Journal of Biogeography 40: 1278–1289.

Thompson JD, Lumaret R. 1992. The evolutionary dynamics of polyploid plants: origins, establishment and persistence. Trends in Ecology & Evolution 7: 302–307.

Tossi VE, Martínez Tosar LJ, Laino LE, Iannicelli J, Regalado JJ, Escandón AS, Baroli I, Causin HF, Pitta-Álvarez SI. 2022. Impact of polyploidy on plant tolerance to abiotic and biotic stresses. Frontiers in Plant Science 13: 869423.

Trøjelsgaard K, Heleno R, Traveset A. 2019. Native and alien flower visitors differ in partner fidelity and network integration. Ecology Letters 22: 1264–1273.

Vamosi JC, Simon GJ, Kennedy BF, Mayberry RJ, Moray CM, Neame LA, Tunbridge ND, Elle E. 2007. Pollination, Floral Display, and the Ecological Correlates of Polyploidy. Functional Ecosystems and Communities 1: 1–9.

Vazquez DP, Lomascolo SB, Belen Maldonado M, Chacoff NP, Dorado J, Stevani EL, Vitale NL. 2012. The strength of plant-pollinator interactions. Ecology 93: 719–725.

Vizentin-Bugoni J, Maruyama PK, de Souza CS, Ollerton J, Rech AR, Sazima M. 2018. Plant-Pollinator Networks in the Tropics: A Review. Ecological Networks in the Tropics: 73–91.

Vollstädt MGR, Galetti M, Kaiser-Bunbury CN, Simmons BI, Gonçalves F, Morales-Pérez AL, Navarro L, Tarazona-Tubens FL, Schubert S, Carlo T, et al. 2022. Plant–frugivore interactions across the Caribbean islands: Modularity, invader complexes and the importance of generalist species. Diversity and Distributions 28: 2361–2374.

Weber MG, Wagner CE, Best RJ, Harmon LJ, Matthews B. 2017. Evolution in a Community Context: On Integrating Ecological Interactions and Macroevolution. Trends in Ecology & Evolution 32: 291–304.

Wei N, Cronn R, Liston A, Ashman TL. 2019. Functional trait divergence and trait plasticity confer polyploid advantage in heterogeneous environments. New Phytologist 221: 2286– 2297.

Wei N, Kaczorowski RL, Arceo-Gómez G, O’Neill EM, Hayes RA, Ashman TL. 2021. Pollinators contribute to the maintenance of flowering plant diversity. Nature 2021 597:7878 597: 688–692.

Wood TE, Takebayashi N, Barker MS, Mayrose I, Greenspoon PB, Rieseberg LH. 2009. The frequency of polyploid speciation in vascular plants. Proceedings of the National Academy of Sciences of the United States of America 106: 13875–13879.

www.inaturalist.org; Nugent J. iNaturalist Citizen science for 21st-century naturalists.

Xiang G, Jiang Y, Lan J, Huang L, Hao L, Liu Z, Xia J. 2023a. Different influences of phylogenetically conserved and independent floral traits on plant functional specialization and pollination network structure. Frontiers in Plant Science 14: 1084995.

Xiang G, Jiang Y, Lan J, Huang L, Hao L, Liu Z, Xia J. 2023b. Different influences of phylogenetically conserved and independent floral traits on plant functional specialization and pollination network structure. Frontiers in Plant Science 14.

Zenil-Ferguson R, Burleigh JG, Freyman WA, Igić B, Mayrose I, Goldberg EE. 2019. Interaction among ploidy, breeding system and lineage diversification. New Phytologist 224: 1252–1265.

