## Supplementary material for "The role of plant polyploidy in the structure of plant-pollinator communities": supplementary_materials.docx

#### Supplementary Notes

#### Note S1: Flower restrictiveness data collection

Flower restrictiveness data were collected for a dataset of 1,434 plant species, with 1,148 of them included in the analysis. Each species was characterized as having phenotypically restrictive or unrestrictive flowers based on flower shape and accessibility of reward (Burns *et al.*, 2019). Thus, flower restrictiveness as defined here broadly reflects the potential for pollinators access to floral rewards. We classified flowers as restrictive if their morphology indicated limited accessibility for a specific group of pollinators to reach floral rewards (e.g.,: bell/funnel, chamber, flag/keel, spur, tubular, gullet, flowers with poricidal anthers, etc.). In contrast, flowers were categorized as unrestrictive if their shape indicated free accessibility to any pollinators (e.g., open/dish, brush or inconspicuous flowers). For members of the Asteraceae family, we considered the capitulum as the baseline for flower shape assessment. Because the capitulum resembles open/dish flowers, all Asteraceae species were considered unrestrictive. Flower restrictiveness information was obtained primarily through species descriptions available in a variety of online sources (e.g., eFloras; GBIF; World Flora Online). When not available, we searched for information in peer-reviewed manuscripts, published datasets (e.g., Bennett et al., 2018; Lanuza et al., 2023), and iNaturalist images (www.inaturalist.org; Nugent, 2018).

#### Note S2: Multivariate analysis to quantify the partial effect of polyploid frequency on network structure

To assess the effect of polyploid frequency on each network index while accounting for the partial contribution of other predictors, we fitted a multivariate spatial autoregressive model (Kissling & Carl, 2008) with the R package spdep (Bivand Roger, 2022) for each network-based index, using all variables as predictors. We also included network size to account for variation in network indices attributed to differences in network size. To avoid collinear predictors, we applied iterative variation inflation factor (VIF) analysis prior to model fitting. Specifically, we iteratively computed the VIF for each predictor and removed those with the highest value until all remaining predictors had VIF values below 2. All plant trait-derived predictors were retained in this process, while among environmental predictors, only BIO10 (temperature at the warmest quarter), BIO15 (precipitation seasonality), and BIO18 (precipitation at the warmest quarter) were kept. The subsequent analyses employed the seven remaining predictors: %PP, %SC, %Restrictive, network size, BIO10, BIO15 and BIO18. The results were consistent with the univariate analysis described in the main text, although some associations that were significant in the univariate analysis became nonsignificant in the multivariate analysis (Table S2).

#### Note S3: Path analysis for modeling the effect of polyploid frequency on network structure – statistical verification.

To assess the robustness of the path analysis, we explored several variations to the initial diagram described in the main text. First, since our base model only accounted for the effect of precipitation on network structure, we examined an alternative diagram where BIO15 (Precipitation seasonality) was replaced with BIO10 (mean temperature of warmest quarter), which was the most influential temperature-based variable in the multivariate analysis (Fig. S3). The results suggested that networks collected from areas with higher temperature are significantly less connected and robust, and that these networks were significantly more enriched with plants having morphologies that restrict pollinator access, and less enriched with self-compatible plants. Importantly, all other effects within the diagram were robust to the replacement of BIO15 with BIO10. Second, we tested whether inclusion of both BIO10 and BIO15 could reveal causal relationships that were not accounted for when considering each class of environmental factor individually (Fig. S4). The inclusion of BIO10 and BIO15 in the diagram simultaneously revealed that the positive effect of precipitation seasonality (BIO15) on modularity is non-significant when accounting for the effect of temperature using BIO10. Similarly, the effect of the frequency of plants with restrictive flowers on modularity became non-significant. However, the remaining effects within the diagram were statistically robust to the inclusion of both environmental factors. Furthermore, the inclusion of BIO10 to the baseline diagram had little contribution to the explanatory power of the model, as evident by the negligible increase in $R^{2}$ values compared to those described in the main text (Fig. 3 and Fig. S4). Third, we examined the separate effects of the frequencies of plants with restrictive flowers and self-compatible plants by fitting the structural equational models while either excluding the %Restrictive factor or %SC factor from the diagram (Fig. S5). This analysis revealed that %SC contributes more to the overall explanatory power of the model compared to %Restrictive, as evident from the consistently higher $R^{2}$values in the %Restrictive-excluded models compared to %SC-excluded ones. The results of the original diagram were statistically robust to the separation of the two factors. These results align with the non-significant effects of %Restrictive on al indices except modularity, compared to the significant effect of %SC on all indices.

#### Note S4: Extinction simulations for modeling the effect of polyploidy on robustness

To examine the explicit effect of polyploid presence within communities on network resilience, we performed extinction simulations using the stochastic coextinction model (SCM) of Vieira & Almeida-Neto (2015). The SCM describes both deterministic and stochastic aspects of coextinctions and can simulate secondary coextinction cascades triggered by primary extinction events. We considered a species extinct when it lost all its interactions in the network.

Here, we used SCM to simulate a complete extinction process by repeatedly selecting a single plant species for primary extinction and triggering a SCM cascade until no interactions remain in the network. Three sets of simulations were generated, each consisting of 100 extinction simulations per network. In the first set, plant species were selected randomly for primary extinction until no interactions remained in the network (termed hereafter ‘random extinction’). In the second set, polyploid species were initially sampled for primary extinction, followed by diploids and then plant species with ploidy unknown ploidy (termed hereafter ‘polyploids-first extinction’). In the third set, diploid species were initially sampled for primary extinction, followed by polyploids and then plant species with unknown ploidy (termed hereafter ‘diploids-first extinction’). Notably, in networks with only few species with available ploidy classification, the constrains on the sequence of species selected for primary extinction were too stringent in both the ‘polyploids-first extinction’ and ‘diploids-first extinction’ scenarios, leading to highly similar simulation outcomes. To mitigate this issue, we limited this analysis to networks containing a minimum of five diploid plants and five polyploid plants, resulting in a dataset of 121 networks.

For each extinction simulation, we computed the robustness index as the area under the extinction curve (Burgos et al. 2007). The robustness score of a network was then computed as the average robustness value across 100 simulations of each scenario. The distributions of mean robustness computed across simulations of each network for the three different scenarios are shown in Fig. S6. The observed distributions were quite similar, with slight left skewness in the "polyploids-first extinction" scenario and slight right skewness for the "diploids-first extinction" scenario. A less variable distribution was obtained for the "random extinction" scenario, possibly because the robustness values across these simulations were less constrained in their selection of species for primary extinction, leading to more balanced mean robustness values per network.

We applied a Wilcoxon test to compare the robustness values computed based on "polyploids-first extinction" and "diploids-first extinction" scenarios. The results consistently showed that the extinction order of plants based on their ploidy level has no significant effect on network robustness (test statistic of 2041 with p-value = 0.7), further indicating that the presence of polyploids has a nonsignificant effect on community robustness, in accordance with our previous analyses. Overall, the results suggest that the effect of polyploid frequency on community robustness is indirect, and as such may be too subtle to be captured via a direct analysis that accounts only for polyploidy while ignoring other community traits.

### Supplementary References

**Bennett JM, Steets JA, Durka W, Vamosi JC, Arceo-Gómez G, Burd M, Burkle LA, Ellis AG, Freitas L, Li J, *et al.*** **2018**. Glopl, a global data base on pollen limitation of plant reproduction. *Sci. Data* **5**: 1–9.

**Bivand Roger**. **2022**. R Packages for Analyzing Spatial Data: A Comparative Case Study with Areal Data.”. **54**: 488–518.

**B. Lanuza J, Rader R, Stavert J, Kendall LK, Saunders ME, Bartomeus I**. **2023**. Covariation among reproductive traits in flowering plants shapes their interactions with pollinators. *Functional Ecology* **37**: 2072–2084.

**Burgos E, Ceva H, Perazzo RPJ, Devoto M, Medan D, Zimmermann M, María Delbue A**. **2007**. Why nestedness in mutualistic networks? *Journal of Theoretical Biology* **249**: 307–313.

**Burns JH, Bennett JM, Li J, Xia J, Arceo-Gómez G, Burd M, Burkle LA, Durka W, Ellis AG, Freitas L, *et al.*** **2019**. Plant traits moderate pollen limitation of introduced and native plants: a phylogenetic meta-analysis of global scale. *New Phytologist* **223**: 2063–2075.

**eFloras**. **2006**. *Taxon*.

**GBIF Occurrence Download**. *(GBIF, 2021)*.

**Kissling WD, Carl G**. **2008**. Spatial autocorrelation and the selection of simultaneous autoregressive models. *Global Ecology and Biogeography* **17**: 59–71.

**Venables WN, B. D. Ripley**. **2002**. *Statistics complements to modern applied statistics with S Fourth edition.*

**Vieira MC, Almeida-Neto M**. **2015**. A simple stochastic model for complex coextinctions in mutualistic networks: robustness decreases with connectance. *Ecology Letters* **18**: 144–152.

**World Flora Online**. *World Flora Online, http://www.worldfloraonline.org/, accessed on August 2023*.

**www.inaturalist.org; Nugent J**. iNaturalist Citizen science for 21st-century naturalists.

### Supplementary Tables

**Table S1.** **Pollination niche of diploids and polyploids.** Higher values of Shannon diversity (calculated from data presented in the original study) reflects pollination generalism and a wider niche breadth, whereas number of pollinator taxa unique to polyploids suggests niche differentiation from diploids.

| Shannon diversity  ***Broader niche in**  **bold** | | Plant taxa | # Plant taxa | # Pollinator taxa (unique to  polyploids) | # Pollinator observations | Calculated from: |
| --- | --- | --- | --- | --- | --- | --- |
| Polyploid | Diploid |  |  |  |  |  |
| **2.38** | 2.17 | *Erythronium* species (4x,2x) | 2 | 14 (7) | 69 | Roccoforte et al.  2015 |
| **2.24** | 1.90 | *Heuchera grosulariifolia* (4x, 2x cytotypes) | 1 | 6 (0) | 1219 | Thompson & Merg 2008 |
| 1.80 | **1.90** | *Chameronium angustifolium* (4x,  2x cytotypes) | 1 | 7 (2) | 167 | Kennedy et al. 2006 |
| 1.45 | **1.62** | *Larrea tridentata* (4x,  2x cytotypes) | 1 | 61 (50) | 1735 | Laport et al. 2021 |

#### Table S2. Results of a univariate regression analysis on all indices, using spatial autoregressive models. For each network-based index, the standardized coefficient ($\boldsymbol{\beta}$) and its LRT-based p-value ($\boldsymbol{p}$) are shown for each examined predictor. Significant coefficients are bolded.

| **Predictor** | **Connectance** | | **Nestedness** | | **Modularity** | | **Robustness** | |
| --- | --- | --- | --- | --- | --- | --- | --- | --- |
|  | $\boldsymbol{\beta}$ | $\boldsymbol{p}$ | $\boldsymbol{\beta}$ | $\boldsymbol{p}$ | $\boldsymbol{\beta}$ | $\boldsymbol{p}$ | $\boldsymbol{\beta}$ | $\boldsymbol{p}$ |
| **%PP** | 0.559 | 0.577 | 1.654 | 0.101 | -0.903 | 0.378 | -0.33 | 0.742 |
| **%SC** | -1.91 | 0.061 | -1.347 | 0.185 | 1.273 | 0.214 | -1.554 | 0.127 |
| **%Restrictive** | -1.374 | 0.17 | -0.208 | 0.835 | **2.786** | **0.005** | -0.793 | 0.429 |
| **BIO4** | **-4.485** | **0** | **-2.138** | **0.028** | 0.802 | 0.421 | **-4.063** | **0** |
| **BIO10** | -1.854 | 0.061 | -0.144 | 0.886 | 1.307 | 0.19 | -0.585 | 0.559 |
| **BIO15** | **-4.66** | **0** | **-2.607** | **0.007** | 0.014 | 0.989 | **-4.099** | **0** |
| **BIO18** | **2.183** | **0.028** | **2.747** | **0.006** | -0.245 | 0.807 | **2.484** | **0.012** |
| **Network size** | **4.83** | **0** | **3.217** | **0.001** | **4.43** | **0** | **3.115** | **0.002** |

**Table S3.** **Results of the multivariate spatial autoregression analysis.** For each network-based index, the standardized coefficient ($\beta$) and its LRT-based p-value ($p$) are shown for the selected predictors. The $R^{2}$ of each model is shown under each response index. Predictors that were not selected for a regression model of a given index are greyed out. Significant coefficients are bolded.

| **Predictor** | **Connectance** | | **Nestedness** | | **Modularity** | | **Robustness** | |
| --- | --- | --- | --- | --- | --- | --- | --- | --- |
|  | $\boldsymbol{R}^{\boldsymbol{2}}$**= 0.37** | | $\boldsymbol{R}^{\boldsymbol{2}}$ **= 0.42** | | $\boldsymbol{R}^{\boldsymbol{2}}$ **= 0.32** | | $\boldsymbol{R}^{\boldsymbol{2}}$ **= 0.29** | |
|  | $\boldsymbol{\beta}$ | $\boldsymbol{p}$ | $\boldsymbol{\beta}$ | $\boldsymbol{p}$ | $\boldsymbol{\beta}$ | $\boldsymbol{p}$ | $\boldsymbol{\beta}$ | $\boldsymbol{p}$ |
| **%PP** | 1.183 | 0.238 | **2.442** | **0.015** | 0.295 | 0.772 | 0.076 | 0.939 |
| **%SC** | **-2.780** | **0.007** | -1.035 | 0.306 | 1.722 | 0.093 | **-2.117** | **0.039** |
| **%Restrictive** | **-1.248** | 0.213 | 0.410 | 0.681 | **2.412** | **0.016** | -1.007 | 0.316 |
| **BIO10** | **-3.156** | **0.001** | -1.518 | 0.129 | 0.241 | 0.809 | -1.360 | 0.176 |
| **BIO15** | **-3.994** | **<**$\boldsymbol{1*}\boldsymbol{10}^{\boldsymbol{-4}}$ | -1.650 | 0.092 | 1.091 | 0.276 | **-3.520** | **0.0002** |
| **BIO18** | 1.750 | 0.08 | **2.565** | **0.011** | -0.673 | 0.500 | 1.693 | 0.087 |
| **Network size** | **4.952** | **<**$\boldsymbol{1*}\boldsymbol{10}^{\boldsymbol{-4}}$ | **2.963** | **0.003** | **4.407** | **<**$\boldsymbol{1*}\boldsymbol{10}^{\boldsymbol{-4}}$ | **2.536** | **0.011** |

### Supplementary Figures

#### Figure S1. The distribution of collected plant-pollinator visitation networks across the globe.

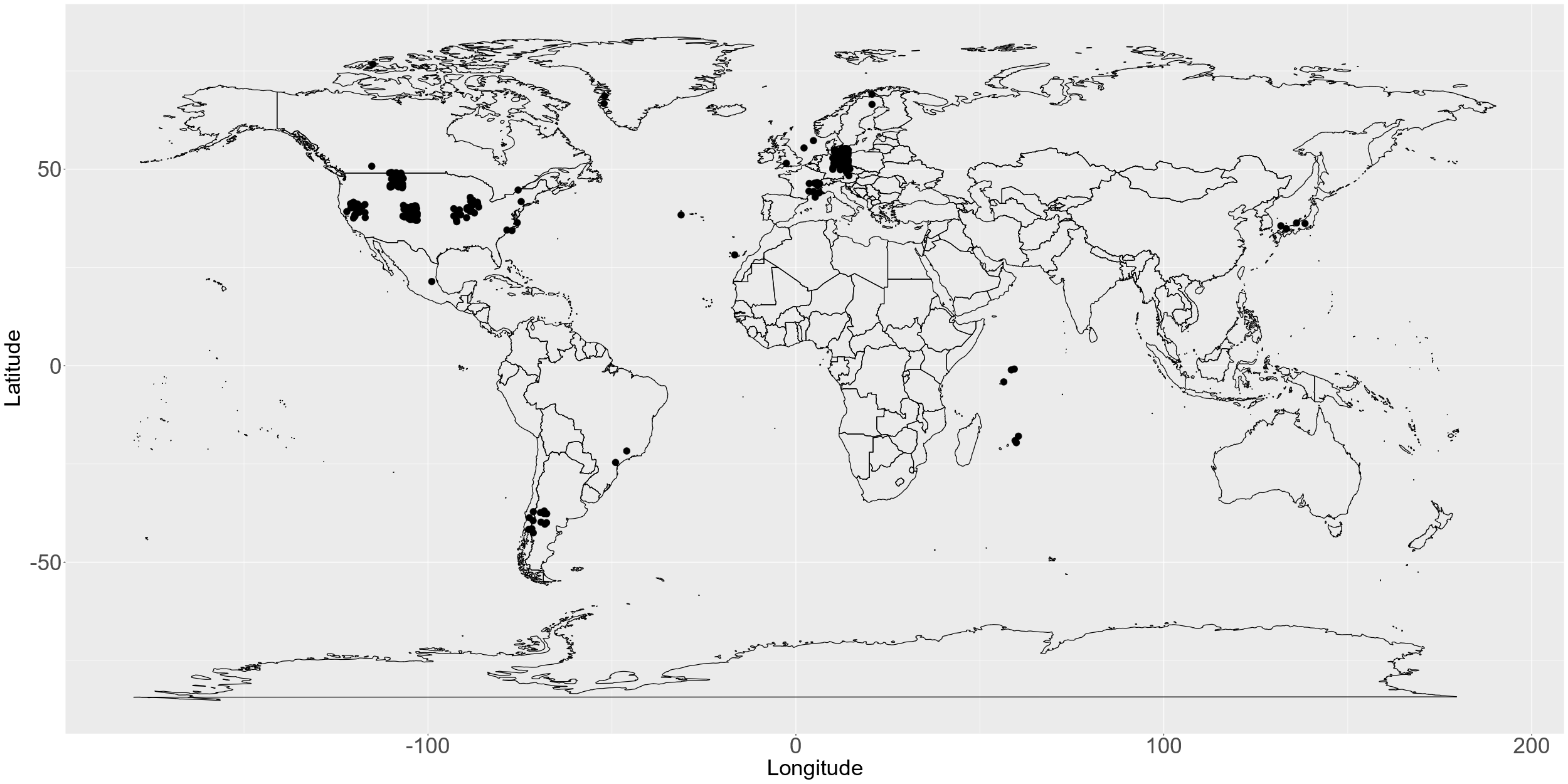

#### Figure S2. The distribution of plant trait data across the analyzed networks. The frequency distribution of polyploids (%PP), plants with flowers that restrict pollinator access (%Restrictive), and self-compatible plants (%SC) are shown across the 316 analyzed networks in panels (a), (b), and (c), respectively. The distributions of missing data for ploidy, flower restrictiveness, and mating system classifications are shown in panels d-f, respectively.

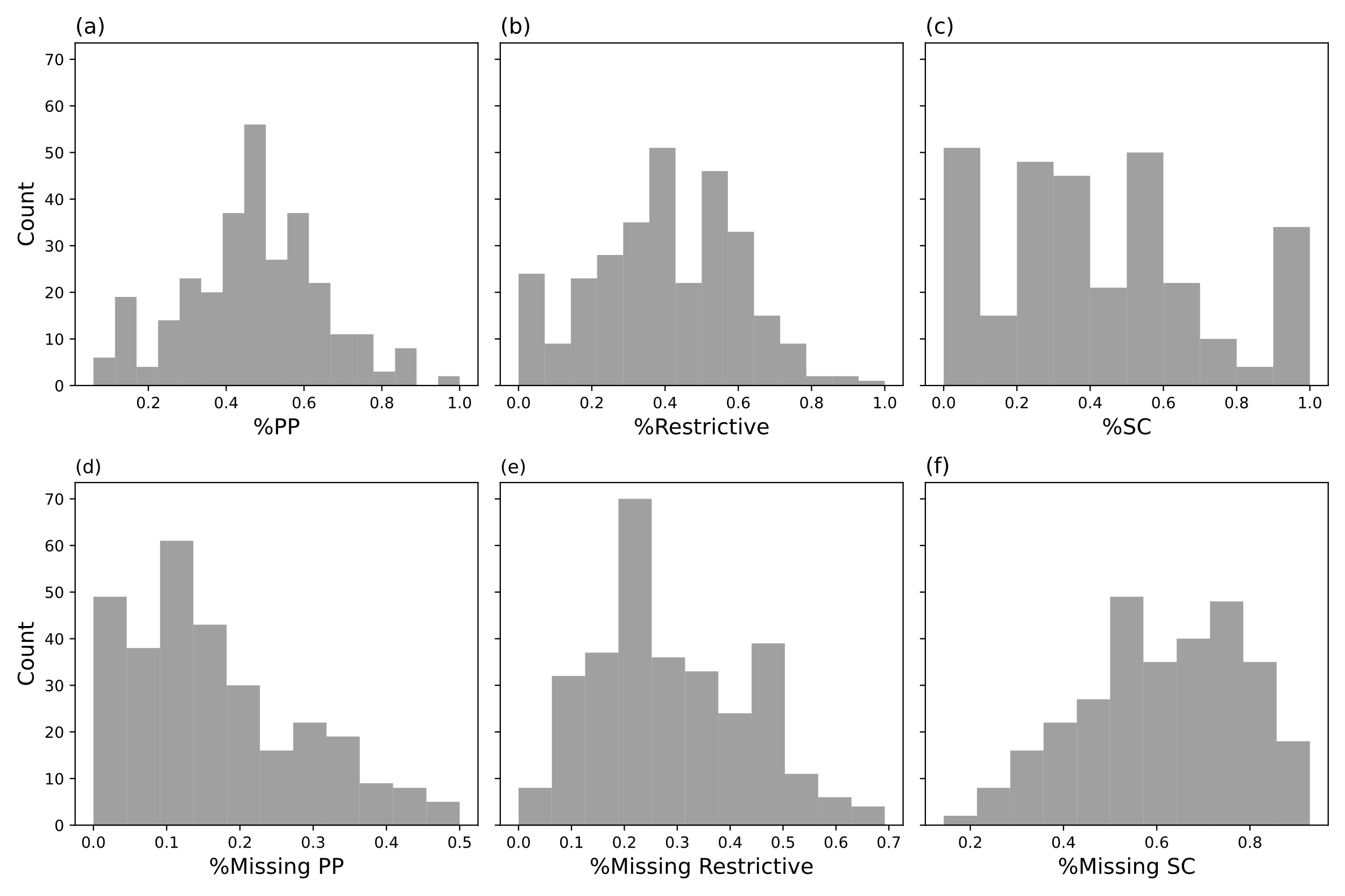

#### Figure S3: Path analysis using temperature (BIO10) as the environmental factor. Path diagrams of the four examined network indices used in the second set of path analysis, in which the environmental factor was BIO10: connectance (a), nestedness (b), modularity (c) and robustness (d). Full lines correspond to paths with significant contribution and dashed lines correspond to paths with nonsignificant contribution. Orange lines correspond to negative coefficients and blue to positive ones. The $\boldsymbol{R}^{\boldsymbol{2}}$ is shown next to each network index. In all panels, the results of the$\boldsymbol{\chi}^{\boldsymbol{2}}$ test for model adequacy were non-significant ($\boldsymbol{p > 0.2}$5), indicating that the model is adequate.

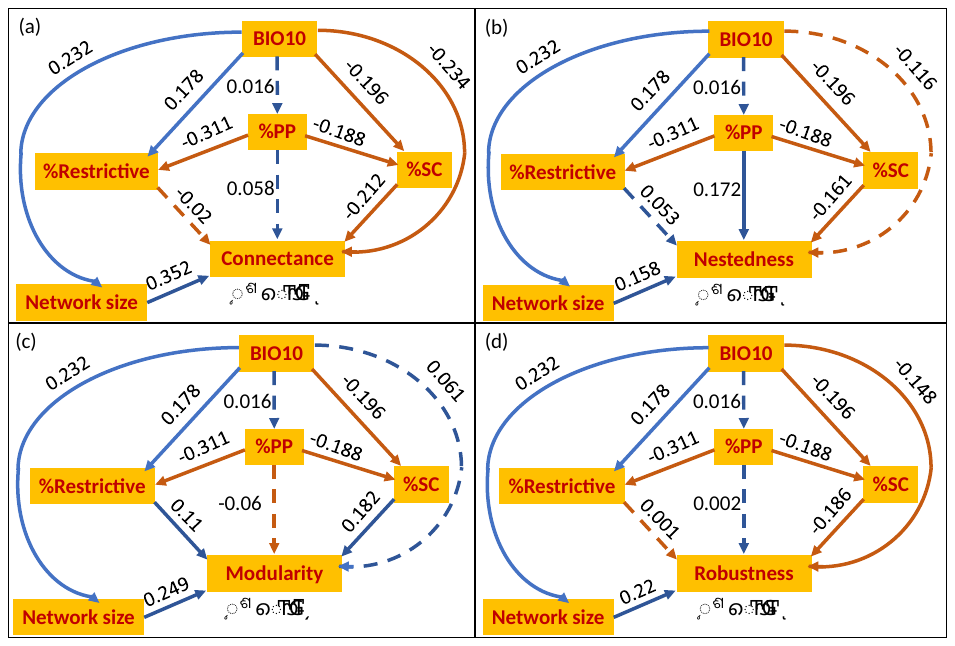

#### Figure S4: Path analysis using two environmental factors. Path Diagrams of the four examined network indices with both BIO10 and BIO15 included as environmental factors: connectance (a), nestedness (b), modularity (c) and robustness (d). Solid lines correspond to paths with significant contribution and dashed lines correspond to paths with nonsignificant contribution. Orange lines correspond to negative coefficients and blue to positive ones. The $\boldsymbol{R}^{\boldsymbol{2}}$ is shown next to each network index. In all panels, the results of the$\boldsymbol{\chi}^{\boldsymbol{2}}$ test for model adequacy were non-significant ($\boldsymbol{p > 0.48}$), indicating that the model is adequate.

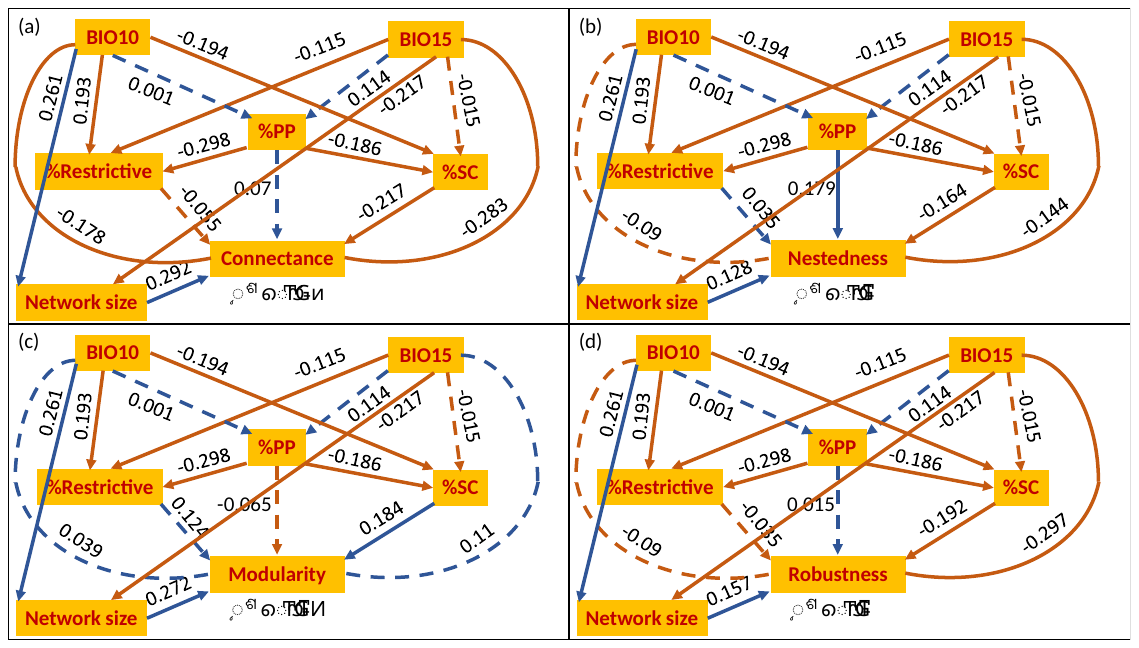

#### Figure S5: Path analysis with selective trait exclusion. Partial diagrams of the four examined network indices: connectance (a,b), nestedness (c,d), modularity (e,f) and robustness (g,h), with %Restrictive excluded (a,c,e,g) or %SC excluded (b,d,f,h). Solid lines correspond to paths with significant contribution and dashed lines correspond to paths with non-significant contribution. Orange lines correspond to negative coefficients and blue to positive ones. The $\boldsymbol{R}^{\boldsymbol{2}}$ is shown next to each network index. In all panels, the results of the$\boldsymbol{\chi}^{\boldsymbol{2}}$ test for model adequacy were non-significant ($\boldsymbol{p>0.08}$ for exclusion of %Restrictive and $\boldsymbol{p>0.16}$ for exclusion of %SC), indicating that the models are adequate.

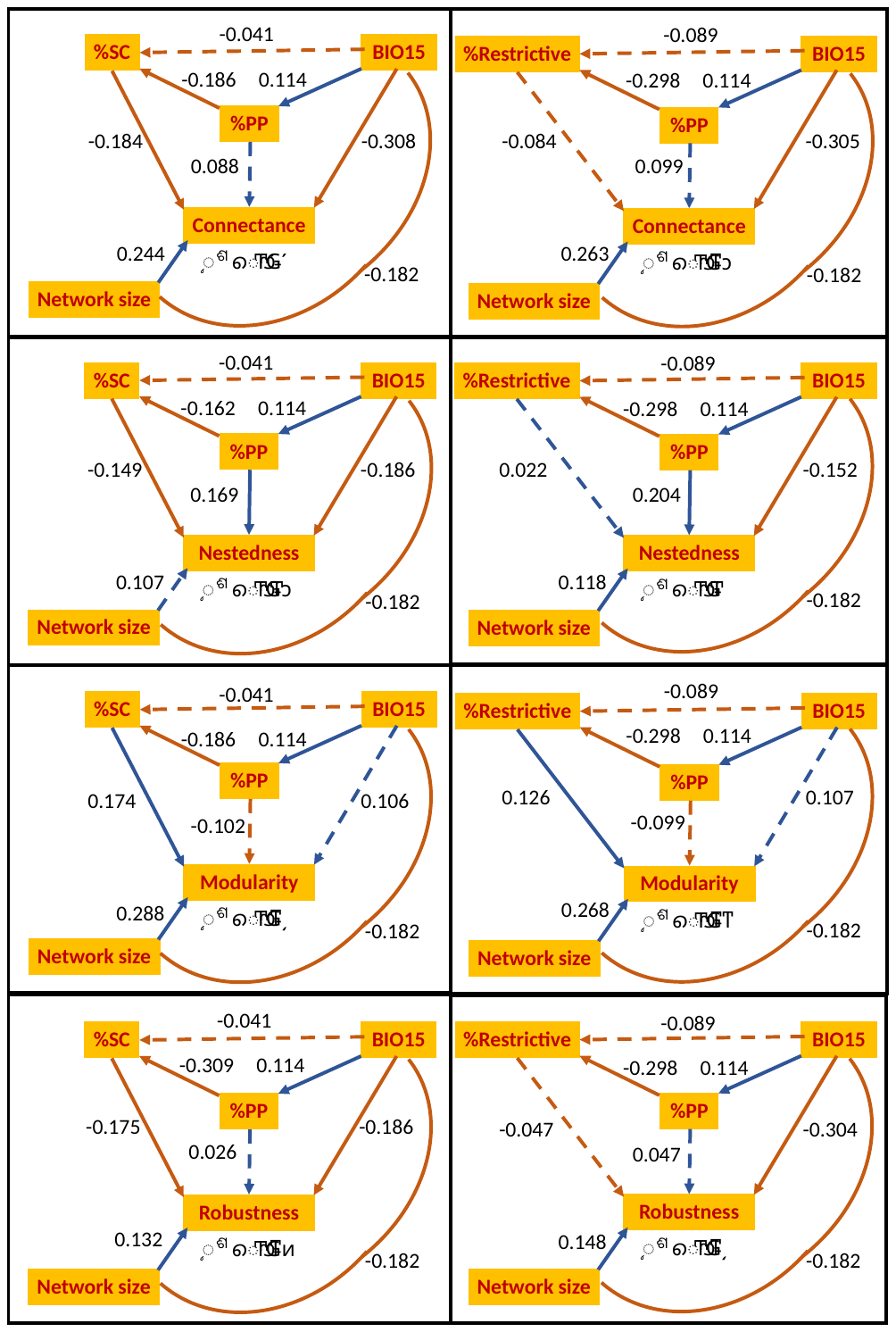

#### Figure S6. Distributions of robustness values under different extinction simulation scenarios. The distribution of mean robustness values across binarized networks is shown, as computed across extinction simulation scenarios with different primary extinction orders: random (grey), polyploids first (orange), and diploids first (blue). The analysis included 121 weighted networks that met the following criteria: at least 50% plant species with available ploidy classification, at least six pollinators, and at least 10 classified plant species (including a minimum of five polyploids and five diploids).

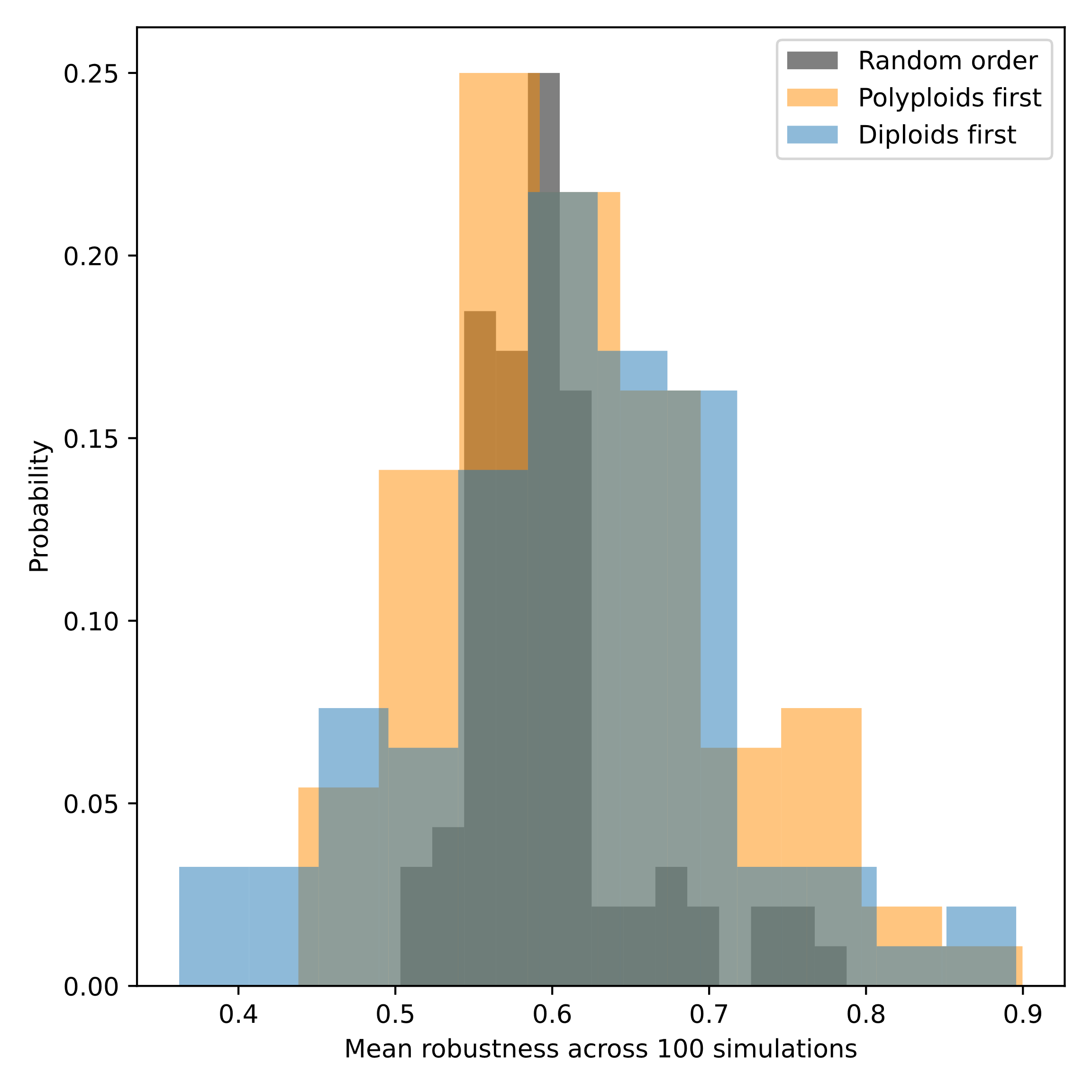
