## Supplementary material for "The role of plant polyploidy in the structure of plant-pollinator communities": supplementary_materials.pdf

**Table S3. Results of the multivariate spatial autoregression analysis.** For each network-based index, the standardized coefficient ($\beta$) and its LRT-based p-value ($p$) are shown for the selected predictors. The $R^2$ of each model is shown under each response index. Predictors that were not selected for a regression model of a given index are greyed out. Significant coefficients are bolded.

| Predictor | Connectance $R^2$= 0.37 | | Nestedness $R^2$ = 0.42 | | Modularity $R^2$ = 0.32 | | Robustness $R^2$ = 0.29 | |
|---|---|---|---|---|---|---|---|---|
| | $\beta$ | $p$ | $\beta$ | $p$ | $\beta$ | $p$ | $\beta$ | $p$ |
| %PP | 1.183 | 0.238 | **2.442** | **0.015** | 0.295 | 0.772 | 0.076 | 0.939 |
| %SC | **-2.780** | **0.007** | -1.035 | 0.306 | 1.722 | 0.093 | **-2.117** | **0.039** |
| %Restrictive | **-1.248** | 0.213 | 0.410 | 0.681 | **2.412** | **0.016** | -1.007 | 0.316 |
| BIO10 | **-3.156** | **0.001** | -1.518 | 0.129 | 0.241 | 0.809 | -1.360 | 0.176 |
| BIO15 | **-3.994** | $<1 * 10^{-4}$ | -1.650 | 0.092 | 1.091 | 0.276 | **-3.520** | **0.0002** |
| BIO18 | 1.750 | 0.08 | **2.565** | **0.011** | -0.673 | 0.500 | 1.693 | 0.087 |
| Network size | **4.952** | $<1 * 10^{-4}$ | **2.963** | **0.003** | **4.407** | $<1 * 10^{-4}$ | **2.536** | **0.011** |

### Supplementary Figures

**Figure S1. The distribution of collected plant-pollinator visitation networks across the globe.**

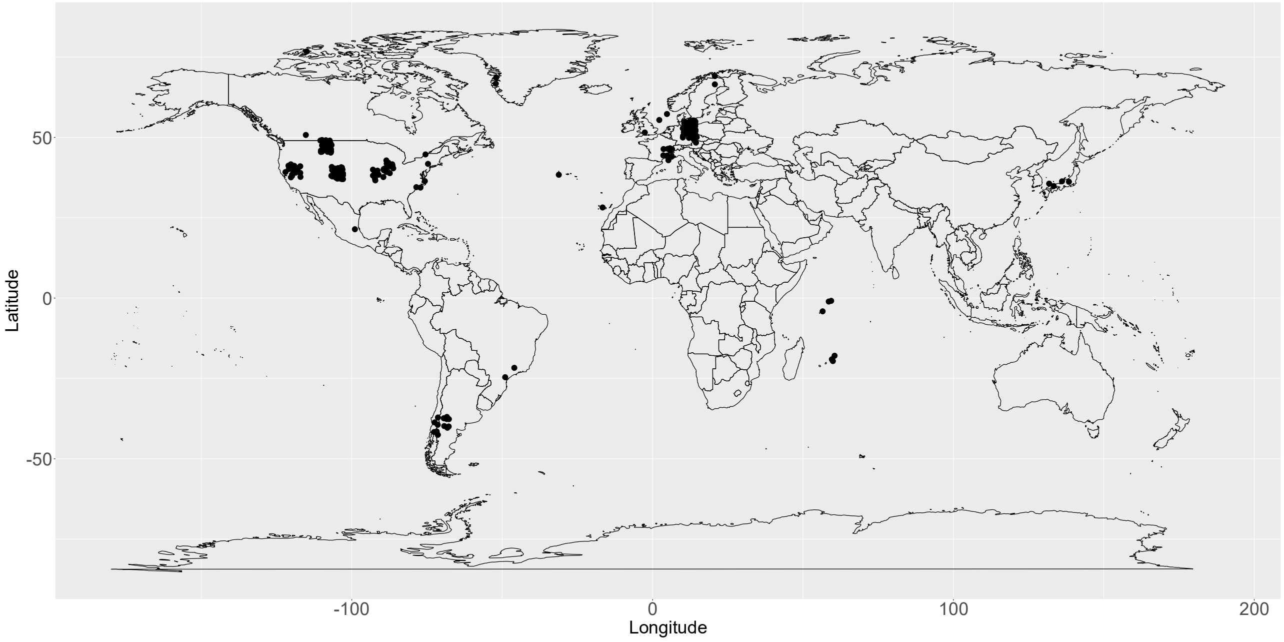

**Figure S2. The distribution of plant trait data across the analyzed networks.** The frequency distribution of polyploids (%PP), plants with flowers that restrict pollinator access (%Restrictive), and self-compatible plants (%SC) are shown across the 316 analyzed networks in panels (a), (b), and (c), respectively. The distributions of missing data for ploidy, flower restrictiveness, and mating system classifications are shown in panels d-f, respectively.

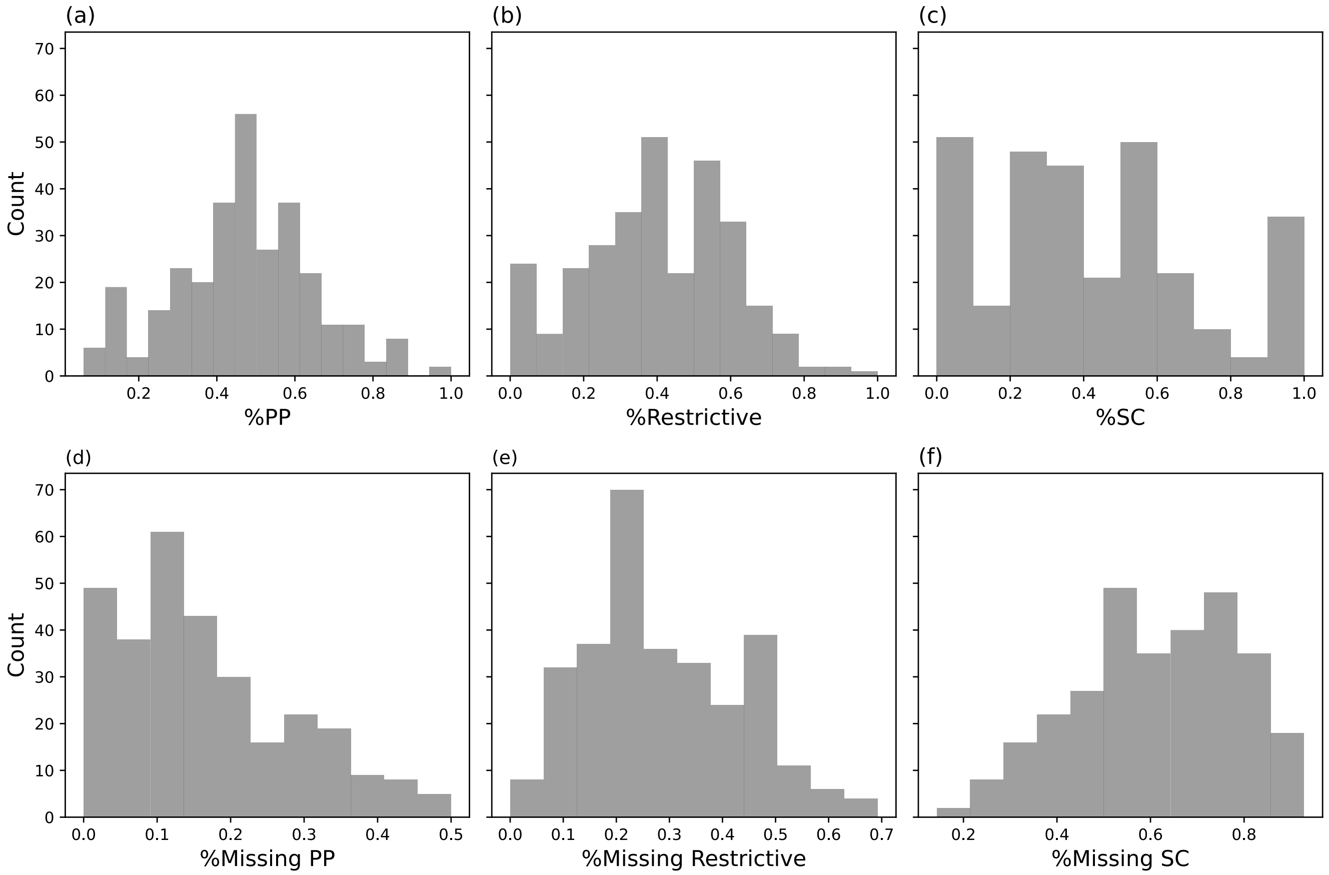

**Figure S3: Path analysis using temperature (BIO10) as the environmental factor.** Path diagrams of the four examined network indices used in the second set of path analysis, in which the environmental factor was BIO10: connectance (a), nestedness (b), modularity (c) and robustness (d). Full lines correspond to paths with significant contribution and dashed lines correspond to paths with nonsignificant contribution. Orange lines correspond to negative coefficients and blue to positive ones. The $\boldsymbol{R^2}$ is shown next to each network index. In all panels, the results of the $\boldsymbol{\chi^2}$ test for model adequacy were non-significant ($\boldsymbol{p > 0.25}$), indicating that the model is adequate.

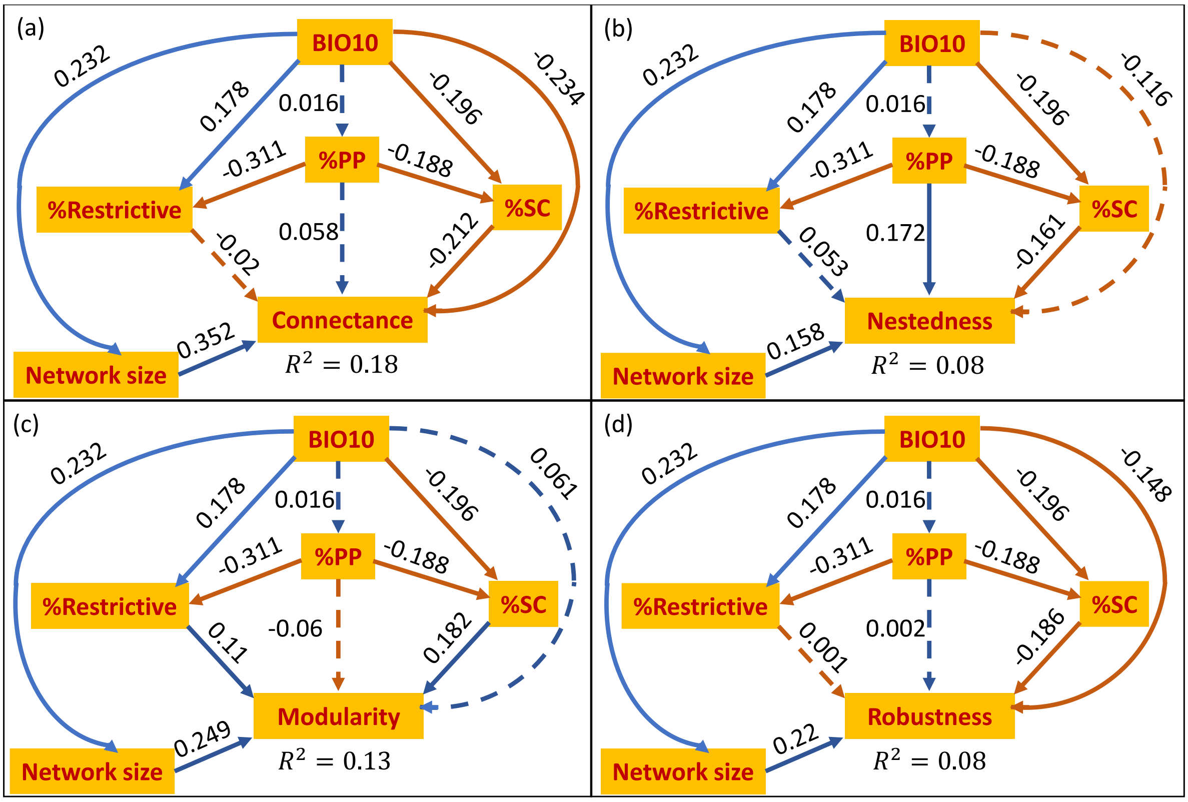

**Figure S4: Path analysis using two environmental factors.** Path Diagrams of the four examined network indices with both BIO10 and BIO15 included as environmental factors: connectance (a), nestedness (b), modularity (c) and robustness (d). Solid lines correspond to paths with significant contribution and dashed lines correspond to paths with nonsignificant contribution. Orange lines correspond to negative coefficients and blue to positive ones. The $\boldsymbol{R^2}$ is shown next to each network index. In all panels, the results of the $\chi^2$ test for model adequacy were non-significant ($\boldsymbol{p} > \mathbf{0.48}$), indicating that the model is adequate.

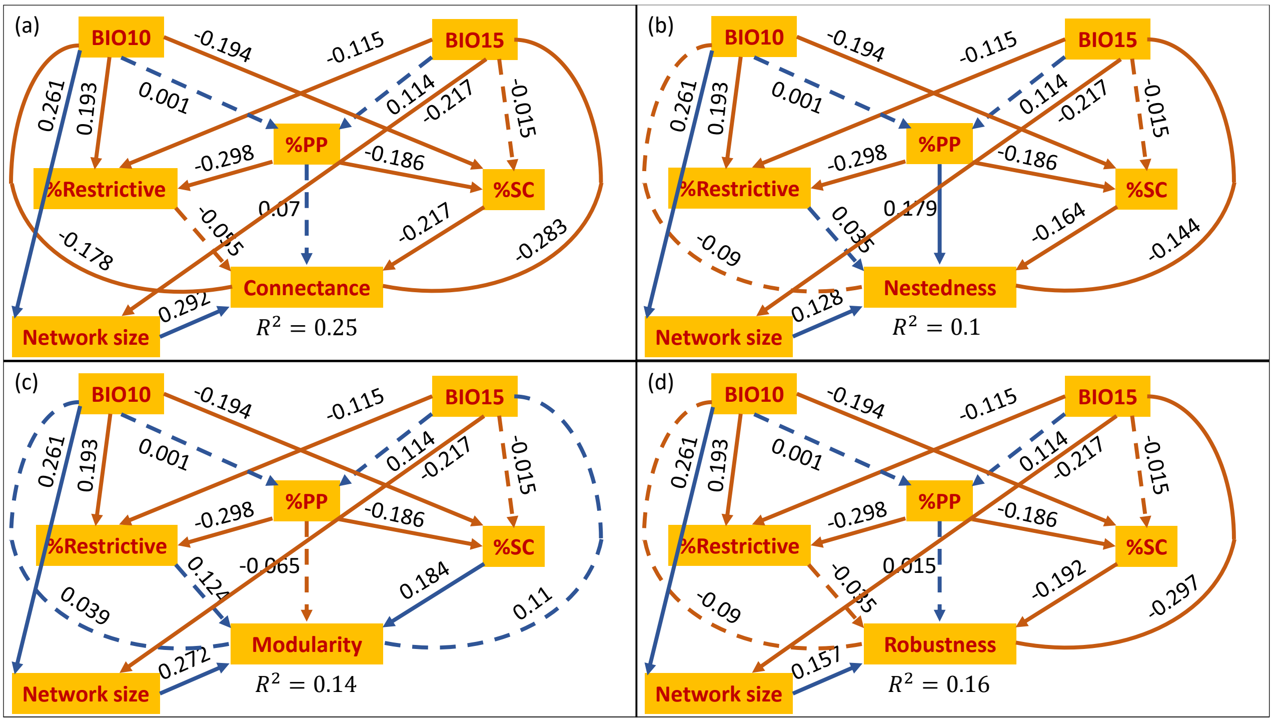

**Figure S5: Path analysis with selective trait exclusion.** Partial diagrams of the four examined network indices: connectance (a,b), nestedness (c,d), modularity (e,f) and robustness (g,h), with %Restrictive excluded (a,c,e,g) or %SC excluded (b,d,f,h). Solid lines correspond to paths with significant contribution and dashed lines correspond to paths with non-significant contribution. Orange lines correspond to negative coefficients and blue to positive ones. The $R^2$ is shown next to each network index. In all panels, the results of the $\chi^2$ test for model adequacy were non-significant ($p > 0.08$ for exclusion of %Restrictive and $p > 0.16$ for exclusion of %SC), indicating that the models are adequate.

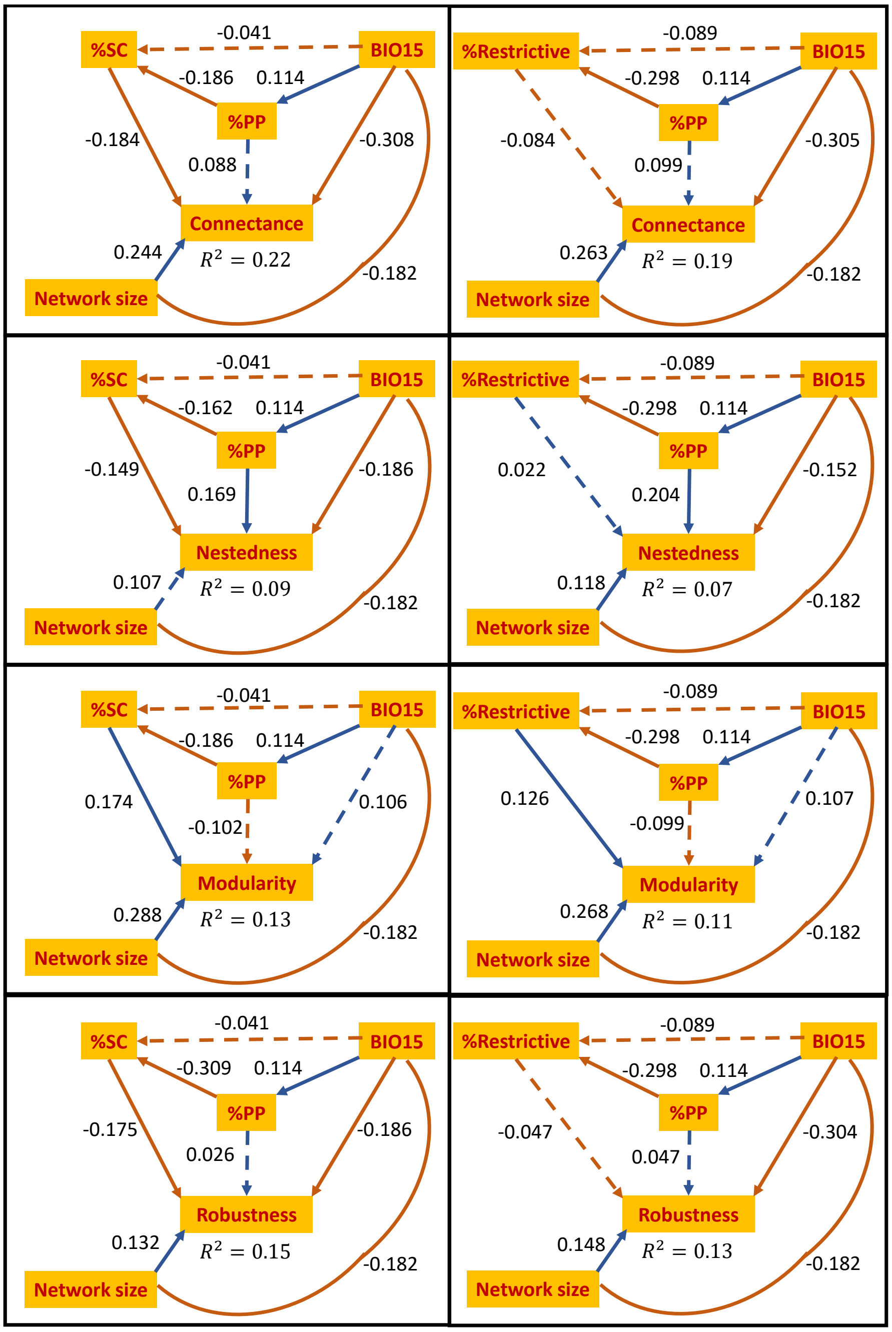

**Figure S6. Distributions of robustness values under different extinction simulation scenarios.** The distribution of mean robustness values across binarized networks is shown, as computed across extinction simulation scenarios with different primary extinction orders: random (grey), polyploids first (orange), and diploids first (blue). The analysis included 121 weighted networks that met the following criteria: at least 50% plant species with available ploidy classification, at least six pollinators, and at least 10 classified plant species (including a minimum of five polyploids and five diploids).

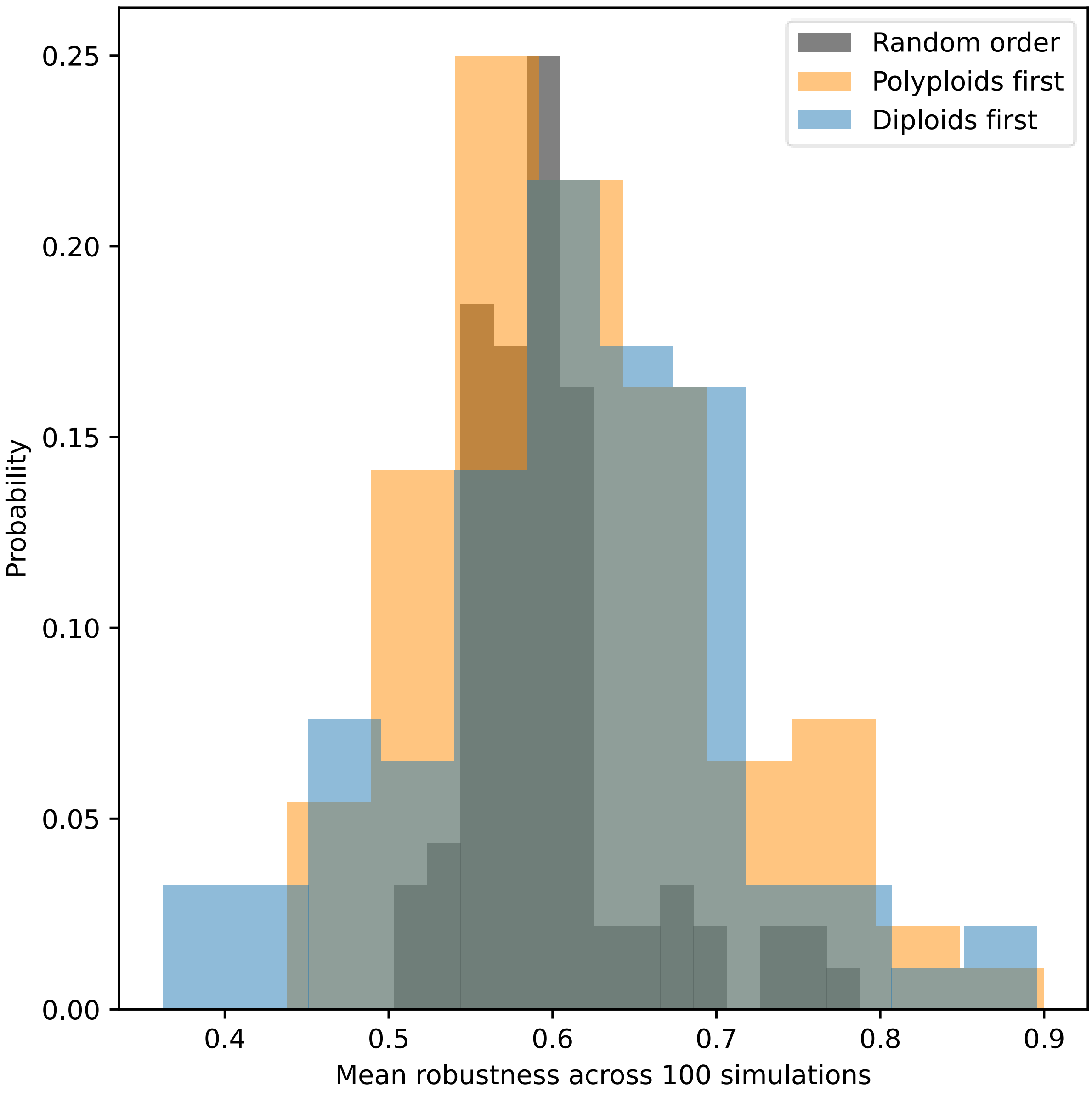
